# Pangenome analyses uncover the scale and evolutionary consequences of structural variation in a wild mammal

**DOI:** 10.64898/2026.01.27.701785

**Authors:** Landen Gozashti, Olivia S. Harringmeyer, Christopher Kirby, Peter Sudmant, Andreas F. Kautt, Hopi E. Hoekstra

## Abstract

Pangenome methods reduce reference bias and reveal genomic variation that is inaccessible with a single reference genome, enabling new insights into genome evolution, particularly in species with high genetic diversity. Here, we constructed a pangenome for the deer mouse (*Peromyscus maniculatus*) – a premier model for studying local adaptation and a potent reservoir of zoonotic pathogens – using long-read assemblies from 14 ecologically and biogeographically distinct populations together with two outgroup species, representing 19 haplotypes. The resulting pangenome spans 3.9 Gb, yet only 1.4 Gb is shared across all deer mouse haplotypes, revealing extensive structural variation within the species. We identified approximately 108.3 million single-nucleotide polymorphisms (SNPs), 23.9 million indels (< 50 bp), and 2.4 million structural variants (> 50 bp), in addition to massive chromosomal rearrangements and an unprecedented diversity of centromeric satellite positions. We also uncovered widespread gene copy-number variation across deer mouse haplotypes, with copy numbers differing by more than an order of magnitude for some genes, and multiple gene-rich loci exhibiting exceptional structural complexity. These genes are enriched for functions related to sensory perception, environmental interaction, and immune responses, and many show signatures of diversifying selection, suggesting that structural variation has contributed to their functional diversification. Together, our results demonstrate the power of pangenome approaches to resolve the full spectrum of genomic variation and reveal a previously underappreciated degree of structural plasticity in mammalian genomes.

## Introduction

Identifying the genetic basis of adaptation and speciation is a central goal of evolutionary biology. Although structural variation has been studied for more than a century, most studies linking genetic variation to phenotypic evolution have focused on single nucleotide variants ^1^. This emphasis may reflect technological limitations rather than evolutionary importance. Until recently, comprehensive discovery of genomic structural variants (> 50 bp in length) was hindered by the limitations of short-read sequencing. Advances in long-read sequencing technologies now enable high-throughput, genome-wide identification of structural variants with unprecedented resolution^1,2^. Comparisons within and between species reveal that structural variants affect several-fold more base pairs than single nucleotide variants – by as much as an order of magnitude ^1–3^ – and often exert larger effects on genome architecture, biological function, and organismal fitness ^1,2^. Despite their abundance and functional importance, the evolutionary origins and consequences of structural variation within species are only beginning to be explored.

Recent advances in pangenome methods have transformed the study of genomic structural variation and enabled a wide range of other genomic applications ^4,5^. Unlike a conventional reference genome, which represents a single, and often collapsed haplotype, a pangenome integrates multiple haplotypes to capture a more complete representation of genomic diversity within a species, thereby reducing reference bias ^4,6,7^. Although early pangenome approaches were largely confined to small prokaryotic genomes, recent developments in bioinformatic methods now support the construction of graph-based pangenomes from multiple large, complex eukaryotic genome assemblies ^8–10^. These graph-based representations not only mitigate reference bias but also enable comprehensive discovery and characterization of complex structural variation ^8,10^, substantially improving the accuracy and sensitivity of structural variant detection relative to reference-based methods ^11,12^. Such advantages should be particularly pronounced in species with high levels of genetic and structural variation ^3,4,6,7^. Yet, despite this promise, pangenome studies have focused largely on humans ^11^, livestock ^13–17^, and agricultural crops ^17–24^, while applications to natural populations –which often harbor substantially more genetic diversity and may therefore stand to benefit the most from a pangenome representations– remain rare ^25^ (but see ^26,27^).

The North American deer mouse (*Peromyscus maniculatus*) provides an exceptional mammalian system for investigating genomic structural variation, local adaptation, and host-pathogen coevolution. As the most abundant mammal in North America, deer mice have adapted to nearly every terrestrial habitat, have large effective population sizes, and have high levels of genetic diversity ^28^. Consistent with this diversity, their genomes exhibit extensive structural variation, including numerous large chromosomal inversions that segregate within natural populations ^29–31^. They also show a recent history of substantial transposable element invasions ^32,33^, which have both actively and passively reshaped genome architecture ^34^. Deer mice are also the principal reservoirs for a wide range of zoonotic pathogens including hantaviruses and the causal agent of Lyme Disease, making them important models for understanding disease ecology, infection tolerance, and host-pathogen coevolution ^35–37^.

To harness this potential, we constructed the first pangenome for a wild mammalian species, integrating 16 deer mouse haplotypes sampled across North America with three haplotypes from two closely related outgroup species. We use this resource to define core and accessory genomic regions, characterize the full spectrum of genomic variation, and identify the genomic features that shape pangenome graph complexity. We further uncover pervasive gene copy-number variation across deer mouse haplotypes suggesting that these regions may serve as hotspots for adaptive gene family evolution. Although purifying selection predominates across most copy-number variable genes, a subset also exhibits signatures of diversifying selection at both the structural and molecular levels, consistent with rapid functional diversification. More broadly, our findings demonstrate the power of pangenome approaches to reveal previously inaccessible genomic variation and establish the deer mouse pangenome as a foundational resource for studies of genome evolution, adaptation, and host-pathogen interactions.

## Results

### High-quality genome assemblies from ecologically diverse deer mouse populations

To maximize the advantages of a pangenome approach, we assembled high-quality *de novo* genomes from genetically and ecologically diverse deer mouse populations spanning much of the species’ North American range (**Figure 1A**). We previously generated chromosome-level assemblies for four populations, including the canonical deer mouse reference population (BW; GCF_049852395.1), using PacBio HiFi long-read sequencing and proximity ligation technologies ^34^. Here, we added nine populations selected to maximize genetic and ecological diversity (see Methods), encompassing forest, prairie, desert, beach, and mountain ecotypes distributed from sea level to elevations exceeding 5,000 feet (**Figure 1A**). Collectively, these populations occupy seven of the thirteen major terrestrial ecoregions inhabited by deer mice, capturing a broad range of environments and the diverse selective pressures associated with local adaptation (**Figure 1A**). To facilitate evolutionary comparisons, we also generated a genome assembly for the closely related white-footed mouse (*Peromyscus leucopus*; ∼9 million years diverged) ^38^. All assemblies were generated using PacBio HiFi sequencing except one, which was generated using Oxford Nanopore data due to limited input material (Supplementary Table 2; see Methods). We additionally generated Omni-C proximity ligation data for five populations to improve scaffolding and haplotype phasing (Extended Data Figure 1; see Methods).

**Figure 1:**
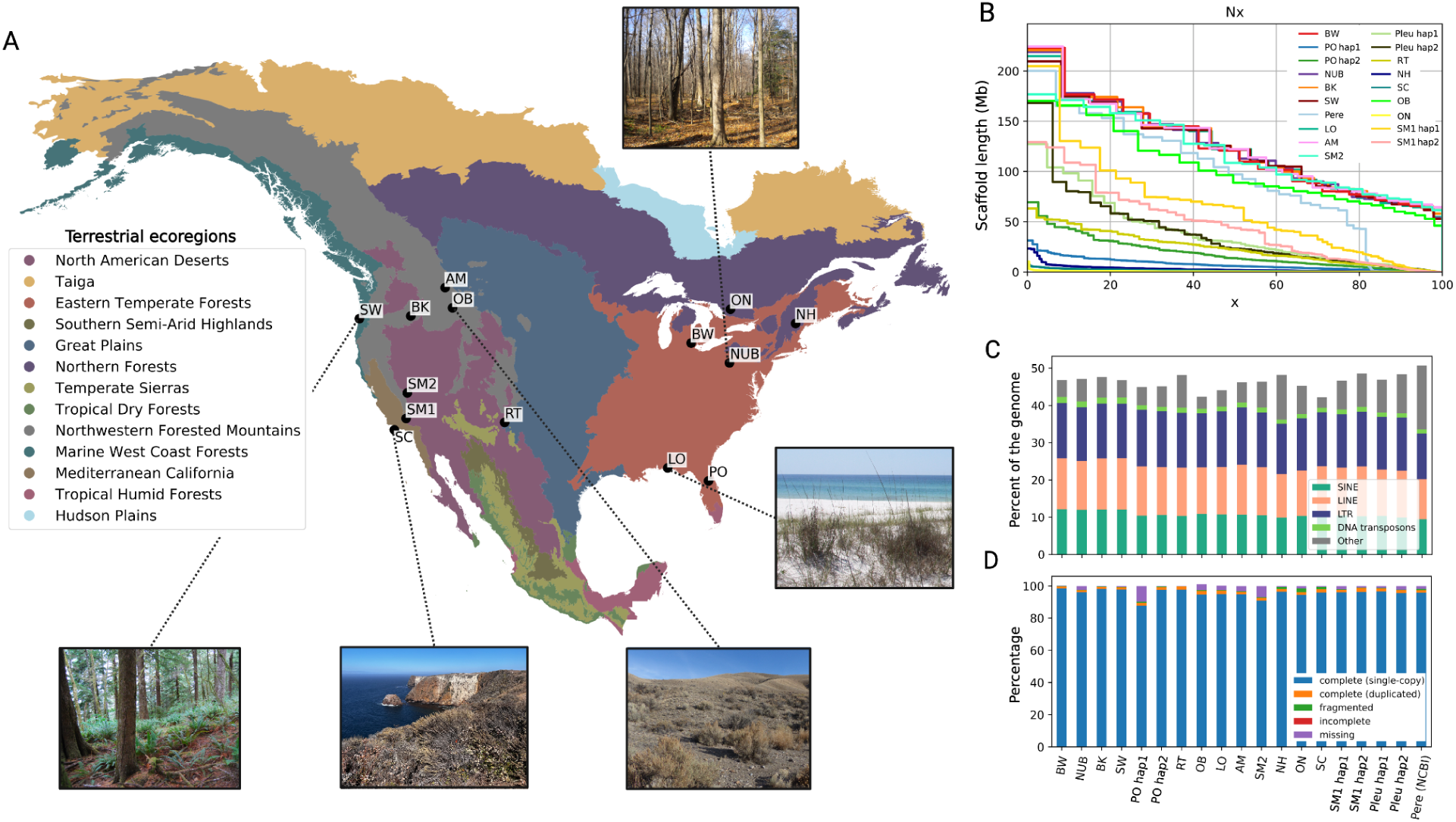
High-quality genome assemblies for ecologically diverse deer mouse populations. (**A)** Map of sampling locations for 14 deer mouse populations selected for pangenome construction (see Table S1 for sample details). Map colors denote level I terrestrial ecoregions as designated by the North American Environmental ATLAS (http://www.cec.org/). Pictures show examples of sampling locations, highlighting habitat diversity of some sampled populations. (Picture credits are as follows: SW = Emily Hager, SC/OB = Andreas Kautt, LO = Nicole Bedford, NUB = Evan Kingsley.) (**B**) Nx plot showing percent of the genome occupied by scaffolds from largest to smallest for each assembly used in the pangenome construction, including 16 haplotypes from 14 populations of *P. maniculatus*, two haplotypes of *P. leucopus* (Pleu), and one haplotype of *P. eremicus* (Pere) (same plot for contigs shown in Extended Data Figure 2; hap1 = haplotype 1, hap2 = haplotype 2). (**C**) Percent of the genome occupied by repeats (SINEs, LINEs, long terminal repeats [LTR], DNA transposons, and other = satellites, simple repeats, and unknown interspersed repeats) in each haplotype assembly. **(D)** Scores across BUSCO categories for all mammalian BUSCO genes in each haplotype assembly.

Overall, the assembled genomes were of high quality despite expected variation in contiguity. Scaffold N50 values ranged from approximately ∼450 kb to ∼122 Mb (contig N50: ∼450 kb - 29 Mb), with a median of 22.4 Mb, while contig counts ranged from 130 to ∼18,600, the latter reflecting the single Oxford Nanopore-based assembly and its distinct assembly pipeline (**Figure 1B**; Extended Data Figure 2; Supplementary Table 2). Lower N50 values for some genomes were anticipated due to DNA degradation observed prior to sequencing (see Methods). We required a minimum mammalian BUSCO completeness score of 89% for inclusion; because five haplotype-phased assemblies did not meet this threshold, we used their corresponding primary assemblies. Consistent with previous observations ^39^, primary assemblies were substantially more contiguous than phased assemblies, with scaffold N50 values up to sixfold higher (**Figure 1B**; Extended Data Figure 2). Overall, included assemblies exhibited BUSCO completeness scores from ∼89.5% to >99% and comparable repeat content, with no evidence that any genome showed substantial underrepresentation of repetitive sequences (**Figure 1C,D**; Supplementary Table 2-3).

### Building deer mouse pangenome graphs

We constructed two complementary deer mouse pangenome graphs using the Pangenome Graph Builder (PGGB) ^10^ and minigraph-cactus (MC) ^9^ **(Figure 2A)**. Both approaches generate nucleotide-resolution graph representations from whole-genome assemblies but differ in how they model genomic variation. PGGB performs all-versus-all genome alignments, including repetitive sequences, to produce a comprehensive and largely reference-independent representation of genomic diversity ^10^. In contrast, MC progressively aligns haplotypes to a single reference backbone, yielding a less complex graph that is computationally efficient for downstream analyses but captures less variation in repetitive regions ^9^. We present both graphs as important resources for the deer mouse model; however, unless otherwise noted, all subsequent analyses are based on the PGGB graph because it provides the most complete representation of structural variation across the genome.

**Figure 2:**
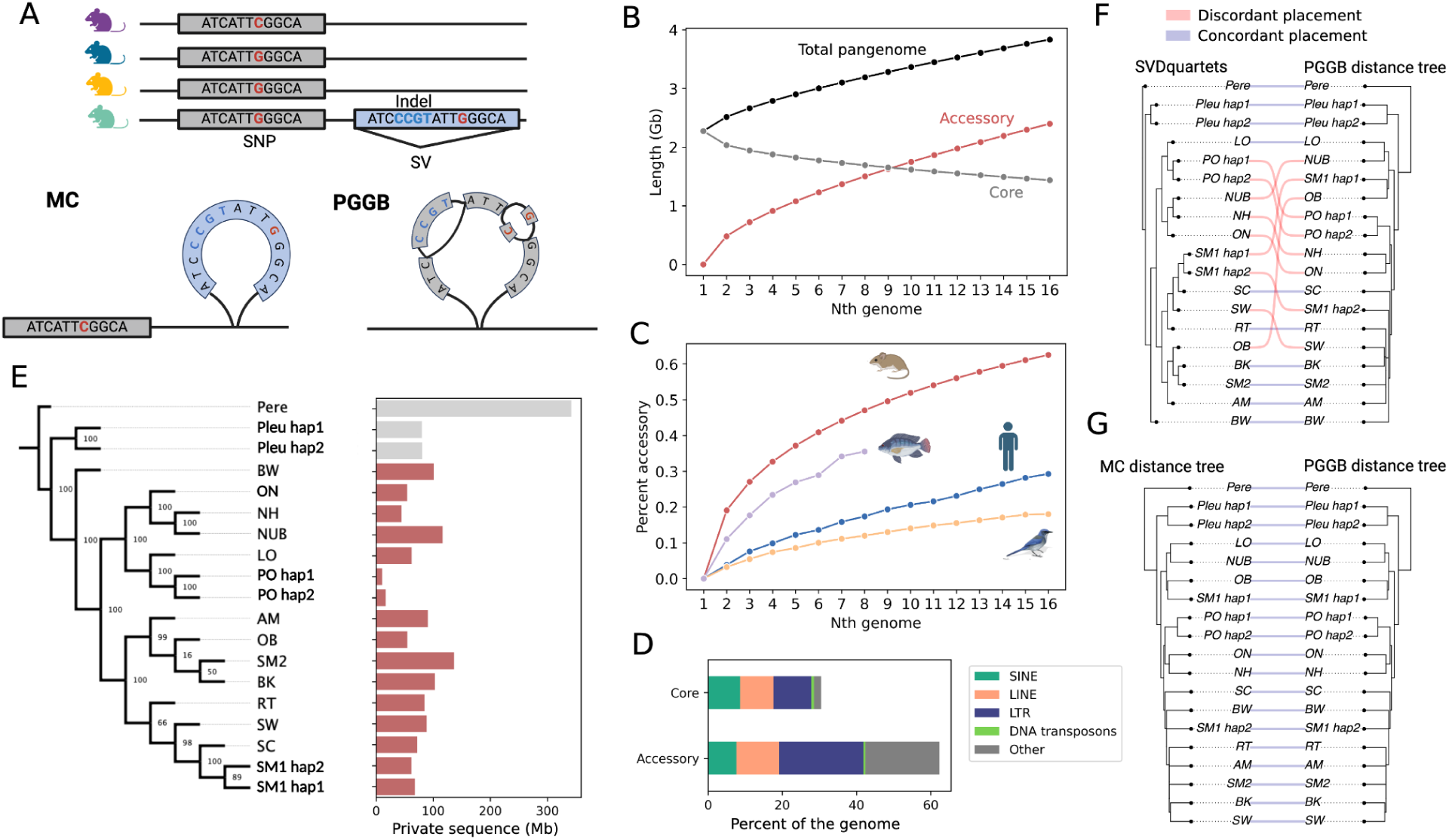
Pangenome reveals extensive accessory genome sequence. **(A)** Schematic representation of MC (minigraph-cactus) and PGGB (Pangenome Graph Builder) pangenome construction methods. MC uses a single reference backbone and progressively aligns additional haplotypes to generate a pangenome graph, whereas PGGB leverages all-versus-all multiple sequence alignments across entire assemblies to generate an unbiased graph in which each variant is represented by a bubble. SNPs are denoted in red, indels in blue, and structural variants by a blue box. (**B**) Growth as a function of increased sampling inferred from the PGGB graph using panacus for the core, accessory, and total pangenome (accessory sequence + core sequence). (**C)** Growth curve inferred by panacus showing the average proportion of the pangenome represented by accessory sequence for the deer mouse PGGB, cichlid minigraph, human PGGB and scrub jay PGGB pangenome (see Methods). (**D**) Percent of the core and accessory genome occupied by different types of genomic repeats (SINE, LINE, LTR, DNA transposons, and other [satellites, simple repeats and unknown repeats]). (**E)** Phylogeny of deer mouse populations and outgroups inferred using SVDquartets from 188,147 unlinked biallelic SNPs and amount of private sequence contributed by each haplotype. Bootstrap support values (N = 100) are displayed above internal nodes. (**F)** Comparison of topology between trees generated by SVDquartets (left) and by Jaccard distances between paths in the PGGB graph (right). (**G**) Comparison of topology between trees constructed from Jaccard distances between paths in the MC graph (left) and the PGGB graph (right).

We constructed chromosome-scale PGGB pangenome graphs from 16 deer mouse genome haplotypes together with three outgroup haplotypes, including the two haplotypes for *P. leucopus* reported here and a previously published long-read-based *P. eremicus* genome, enabling downstream polarization of genomic variants. Following recommendations from the Human Pangenome Reference Consortium, we generated independent graphs for each chromosome because all-versus-all genome alignment does not scale efficiently to interchromosomal comparisons ^11^ (see Methods). This strategy is well-suited to deer mice, which exhibit little evidence of interchromosomal rearrangement from both cytogenetic and genomic analyses ^34,40,41^. We assigned assembly contigs to chromosomes based on sequence similarity (see Methods), recovering 23 autosomal clusters and one X chromosome cluster (Extended Data Figure 3A). A small set of highly repetitive, ambiguously assigned contigs (mean 85% repeat content compared to 45% in clustered) was excluded from downstream analyses, as was the X chromosome, because phased assemblies were unavailable for male samples. The resulting autosomal pangenome comprises 832,281,886 nodes, each DNA sequence shared by one or more haplotypes, connected by 1,196,418,734 edges that record which segments are adjacent in the haplotypes (Supplementary Table 4). Graph complexity varied substantially among chromosomes and increased with repeat content, indicating that repetitive DNA is a major determinant of pangenome graph structure (Extended Data Figure 3B; Kendall’s Tau = 0.38, P = 0.01).

### Underappreciated genetic diversity and accessory sequence within deer mice

Most mammalian model organisms harbor relatively low levels of genetic diversity, leaving the full extent of genetic variation in natural mammalian populations largely unexplored. To quantify genome sharing among deer mice, we compared the core genome (sequences present in all haplotypes) with the accessory genome (sequences absent from at least one haplotype) after excluding outgroup assemblies. Strikingly, the deer mouse pangenome contains a remarkable 2.5 Gb of accessory sequence but only 1.4 Gb of core sequence, meaning that approximately 60% of all sequence is accessory (**Figure 2B**; Extended Data Figure 4). This proportion greatly exceeds that reported for other vertebrate pangenomes, including in humans (28%), scrub-jays (18%), and multiple species of cichlid fishes (35%) (**Figure 2C**; see Methods). As in other species, accessory sequences are highly enriched for repetitive elements, whereas the core genome is comparatively repeat-depleted (**Figure 2D**; Supplementary Table 5).

The extensive accessory genome highlights the limitations of a single linear reference genome. Approximately 1.25 Gb of sequence in the pangenome is absent from the traditional deer mouse “reference genome” (BW). More generally, any individual haplotype would omit an average of 1.55 Gb (∼40%) of pangenome sequence if used as the sole reference, indicating that no single genome adequately represents species-wide diversity (Extended Data Figure 5). Each deer mouse haplotype also contributes substantial private sequence, averaging 73 Mb (range = 10–139 Mb; **Figure 2E**). Remarkably, several deer mouse haplotypes contain more private sequence than the *P. leucopus* outgroup, indicating that structural differences among deer mouse populations can rival those observed between species.

This abundance of private and broader accessory sequences likely reflects numerous independent structural variants segregating among populations. Consistent with this observation, phylogenetic relationships inferred from pangenome graph structure differ markedly from those reconstructed using 188,147 unlinked biallelic SNPs (normalized Robinson-Foulds distance > 0.8; **Figure 2F**; Extended Data Figure 6), and the amount of private sequence is positively correlated with phylogenetic discordance (Kendall’s Tau = 0.400, P=0.045). Importantly, PGGB- and MC-derived graphs produced identical topologies (normalized Robinson-Foulds distance = 0; **Figure 2G**). Together, these results reveal an unexpectedly large accessory genome and an exceptional degree of structural genomic diversity across natural deer mouse populations.

### Pangenome decomposition reveals widespread structural variation

Deer mouse genomic variation is shaped more by structural variation than by single nucleotide variation. Pangenome graph decomposition revealed extensive genetic variation across deer mouse haplotypes. In total, we identified 108,259,150 SNPs, 23,915,976 indels (< 50 bp), and 2,429,347 structural variants (> 50 bp), collectively affecting approximately 4.7%, 5.6% and 29.5% of callable genomic sequence, respectively (**Figure 3A,B**; see Methods). Although less numerous than SNPs, structural variants span more than six times as much of the genome, indicating that structural mutations have likely contributed disproportionately to genome evolution in deer mice.

**Figure 3:**
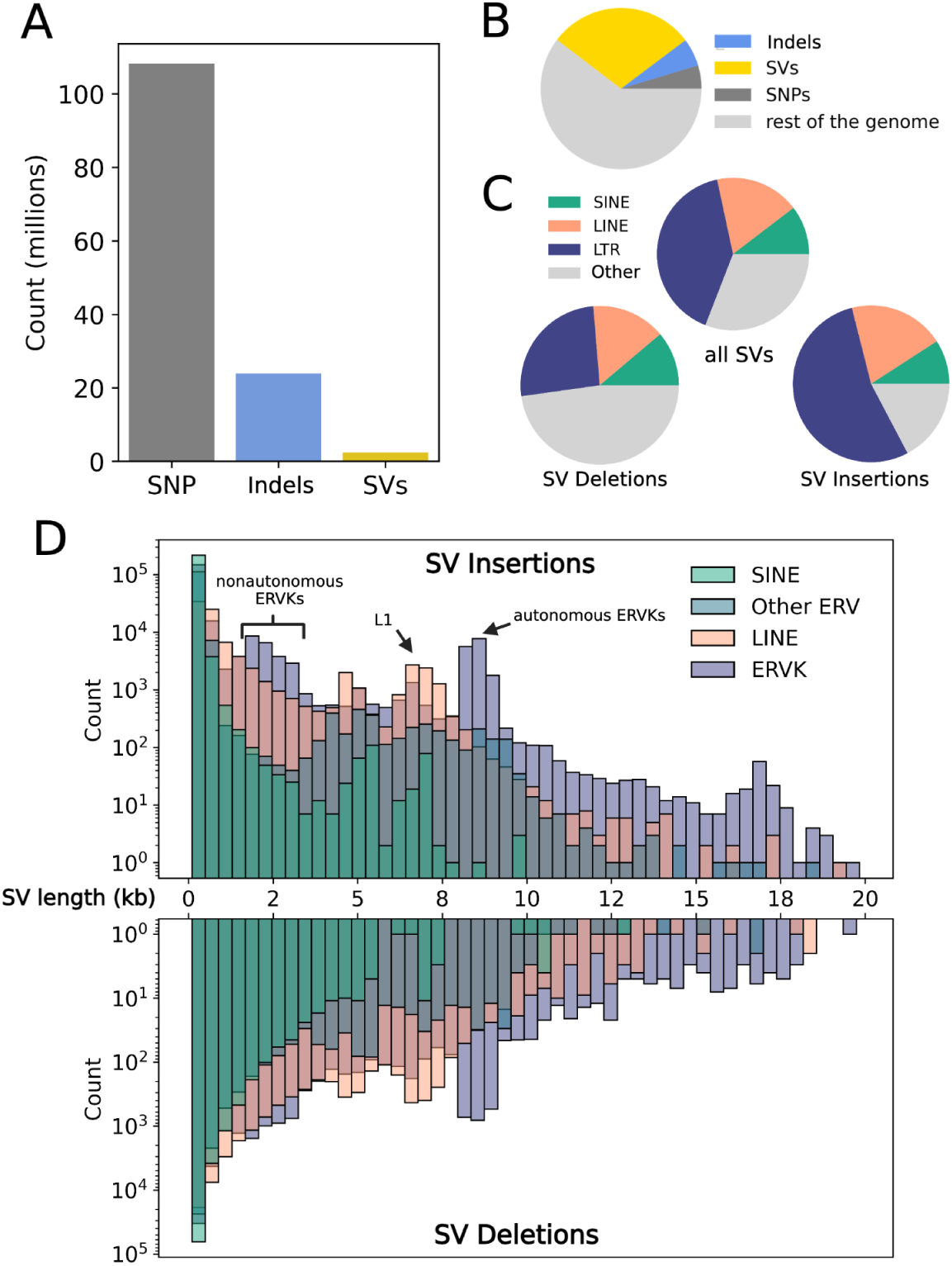
Impact of structural variants and TEs on genomic variation. **(A)** Counts for SNPs, indels (<50 bp) and SVs (>50 bp) identified by pangenome decomposition. (**B**) Proportion of the genome affected by variants across variant classes (SNPs, indels, SVs). (**C)** Proportion of sequence occupied by transposable elements for all SVs, polarized deletions, and polarized insertions. (**D)** SV length distributions for polarized SV insertions and deletions associated with specific repeats (SINE, LINE, ERVK, other ERV). Distributions show distinct peaks (arrows) corresponding to the lengths of active transposable elements as well as non-autonomous derivatives ^32^.

To infer the evolutionary origins of these variants, we compared each variant with the outgroup species, *P. eremicus* and *P. leucopus*, allowing the majority of biallelic indels and structural variants (SVs) to be polarized as either insertions or deletions. This analysis identified 9,940,910 indel insertions, 12,962,871 indel deletions, 1,295,624 SV insertions, and 1,005,962 SV deletions. We also detected 1,244,601 complex indels and SVs with alleles of different lengths that could not be unambiguously polarized as simple insertions or deletions, together with a smaller set of multiallelic and ambiguously polarized variants (Extended Data Figure 7). These results demonstrate that structural variation in deer mice encompasses a broad spectrum of mutational classes extending well beyond canonical insertions and deletions.

### Endogenous retroviruses account for much of the structurally variable sequence

Retrotransposons account for the majority of structural variation across deer mouse populations. Although transposable elements (TEs) occupy approximately 40% of the genome, they comprise nearly 69% of all structurally variable sequence, representing a highly significant enrichment relative to genome-wide expectations (**Figure 3C**; chi-square test, P<0.0001). This enrichment is driven primarily by endogenous retroviruses (ERVs), which account for ∼41% of structurally variable sequence despite comprising only ∼15% of the genome (**Figure 1C**; **Figure 3C**).

Structural variants also differ markedly in TE composition across mutational classes. TEs comprise approximately ∼83% of insertion sequences but only ∼52% of deletion sequences (chi-square test, P<0.0001), consistent with recent and ongoing TE accumulation in deer mouse populations (**Figure 3C**). Using a curated deer mouse TE library (see Methods) ^32^, we identified polymorphic TE insertions underlying ∼80% of SV insertions and 27% of SV deletions. Although SVs range in size from 50 bp to hundreds of kilobases, more than 99% are shorter than 20 kb, and their size distributions exhibit distinct peaks corresponding to recently active retrotransposon families, including LINE1s (L1) as well as autonomous and nonautonomous ERVKs (ERVs of betaretroviral origin in deer mice ^32^) (**Figure 3D**). Previous studies have reported circumstantial evidence of a massive invasion of ERVK elements, but evidence for ongoing invasions at the population level was previously lacking ^32,33^. Polymorphic TEs also show very low Kimura divergence from their consensus sequence (0-5%), providing additional evidence for recent transpositional activity (Extended Data Figure 8). Together, these findings identify recent ERV-driven TE invasion as the dominant force generating structural variation in contemporary deer mouse genomes.

### Pangenome graphs illuminate complex chromosome-scale structural variation

Pangenome graphs reveal complex, chromosome-scale patterns of genetic variation that are difficult to resolve using linear genome representations. Every deer mouse chromosome contains regions of exceptionally high graph depth, indicating that multiple haplotypes repeatedly traverse the same graph nodes (**Figure 4A**). Two-dimensional graph visualization shows that these regions form dense clusters of interconnected bubbles, reflecting highly complex sequence architectures (**Figure 4B**). We identified high-depth regions (≥100 kb with average node depth >500 ^26^) on every chromosome. Most chromosomes contain multiple high-depth regions (median = 5; **Figure 4C**), indicating that complex repetitive structures are broadly distributed throughout the genome. An unusually large number of 220 high-depth regions on chromosome 1 is mostly attributable to megabase-scale hotpots of endogenous retroviruses, segmental duplications, and highly repetitive KRAB zinc-finger (KZNF) genes ^32^ (Extended Data Figure 9). The origins of high-depth regions on the remaining chromosomes are less apparent, so we tested whether they were enriched for specific repeat classes.

**Figure 4:**
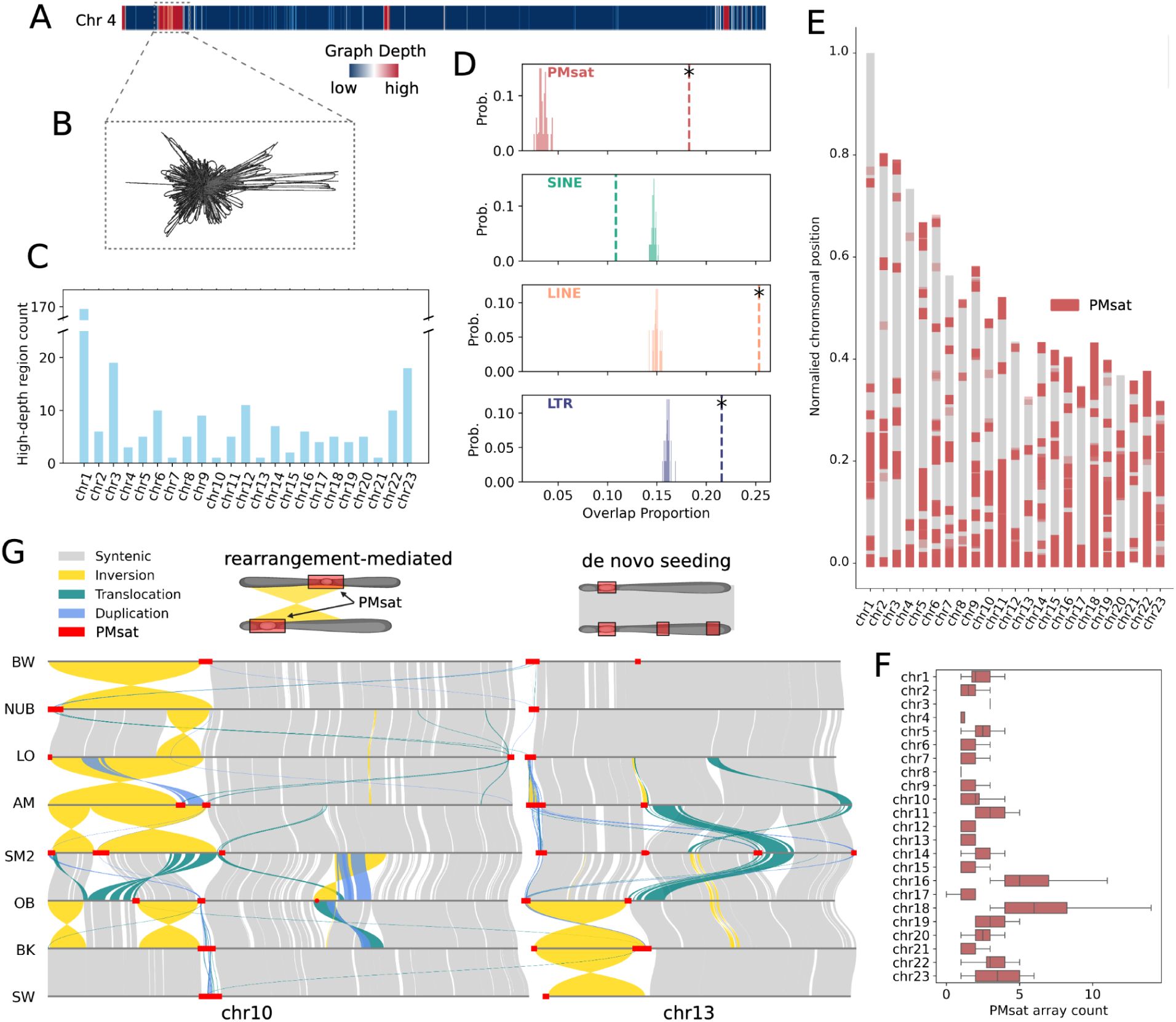
Complex chromosome-scale structural variation. **(A**) Compressed graph depth across the chromosome 4 PGGB graph (blue = low graph depth; red = high graph depth). (**B**) Example of 2-dimensional graph layout for a high-depth region on chromosome 4. High-depth regions are defined as contiguous regions with average node depth > 500 (see Methods). (**C**) Counts for high-depth regions for each chromosome. (**D**) Expected overlap between high-depth regions and different repeats from 1,000 random resampled permutations (histograms) compared to observed overlap (dotted lines). Asterisks indicate significant enrichment (FDR-corrected two-sided permutation test; Q < 0.001). (**E**) Normalized position of PMsat arrays (red) across all haplotypes for each chromosome. PMsat array positions are plotted with alpha = 0.7 such that regions of higher PMsat density appear darker and regions with lower PMsat density appear lighter. (**F**) Distribution of interspersed PMsat array counts across haplotypes for each chromosome. Box center lines show the median. Lower and upper bounds for boxes represent the first and third quartile, respectively. Whiskers extend to the minimum and maximum values within 1.5× the interquartile range. (**G**) Synteny plot for chromosomes 10 and 13 of assemblies scaffolded with Omni-C data. Syntenic regions, inversions, duplications, and translocations are shown in gray, yellow, purple, and blue, respectively; PMsat arrays are shown as red bars along chromosomes. On chromosomes such as chr10 (left), PMsat array repositioning is associated with large chromosomal inversions. Other chromosomes such as chr13 (right) exhibit interspersed PMsat arrays in the absence of chromosomal rearrangements.

Previous studies have shown that high-depth regions frequently correspond to satellite-rich loci such as centromeres or telomeres ^26,42^. We therefore tested whether high-depth regions in deer mice were enriched for specific repeat classes, including the conserved centromeric satellite repeat PMsat. High-depth regions showed a striking seven-fold enrichment for PMsat, together with more modest but significant enrichments for recently active TEs such as LTR and LINE retrotransposons (**Figure 4D**; FDR corrected permutation test, N=1000, Q<0.001). These findings support the hypothesis that dispersed PMsat arrays are a major source of graph complexity and are consistent with previously proposed models of dynamic centromere position in deer mice ^34,43,44^. More broadly, they demonstrate that the topology of the pangenome graph can identify large-scale repetitive genomic features that are difficult to detect with conventional reference-based approaches.

To characterize the diversity of centromere positions across deer mouse haplotypes, we annotated PMsat arrays in each haplotype using the previously identified deer mouse centromeric satellite sequence. Comparison of PMsat locations reveal an unprecedented diversity of centromere positions across the pangenome. Every autosome exhibited variation in centromeric satellite position or presence/absence variation (**Figure 4E,F)**. On several chromosomes, including chromosome 18 and chromosome 23, the cumulative span of PMsat arrays across all haplotypes exceeds that of non-PMsat sequence (**Figure 4E)**, highlighting the remarkable extent of centromere positional variation captured within the pangenome.

Previous cytological and genomic studies have documented frequent centromere repositioning in deer mice, but the underlying genomic mechanisms remain unclear ^34,43,44^. To distinguish among possible mechanisms, we compared Hi-C-supported chromosome assemblies using Syri to identify syntenic regions and large-scale chromosomal rearrangements, including inversions, duplications, and translocations. Some PMsat repositioning events were clearly associated with large chromosomal inversions. For example, on chromosome 10, multi-megabase inversions consistently shift PMsat arrays between central and distal chromosomal positions (**Figure 4G**). In contrast, chromosome 13 contains numerous PMsat presence-absence polymorphisms despite extensive collinearity (**Figure 4G**), indicating that centromeric satellite arrays can also arise or be lost independently of large-scale rearrangements. Although recurrent inversions cannot be excluded ^45^, these observations suggest that multiple mechanisms contribute to the remarkable diversity of centromere positions across deer mouse chromosomes.

### Widespread gene copy number variation across deer mouse populations

Our assembly-based pangenome provides a unique opportunity to characterize gene copy number variation (CNV) and haplotype diversity across natural mammalian populations. To do so, we constructed a pangene graph, in which nodes represent genes and edges capture their adjacency and orientation within each haplotype, enabling robust identification of gene copy number and haplotype variation while minimizing the complexity introduced by repetitive intergenic sequence and collapsed paralogs ^46^. Among the 20,956 autosomal protein-coding genes in the BW reference genome that could be confidently evaluated, 6,158 (∼29%) exhibit copy number variation across deer mouse populations (**Figure 5A**; Supplementary Table 6). This far exceeds the proportion of copy number variable genes reported in humans (∼5%) ^47^, indicating that CNVs represent a pervasive and likely underappreciated source of genetic diversity in wild mammalian populations.

**Figure 5:**
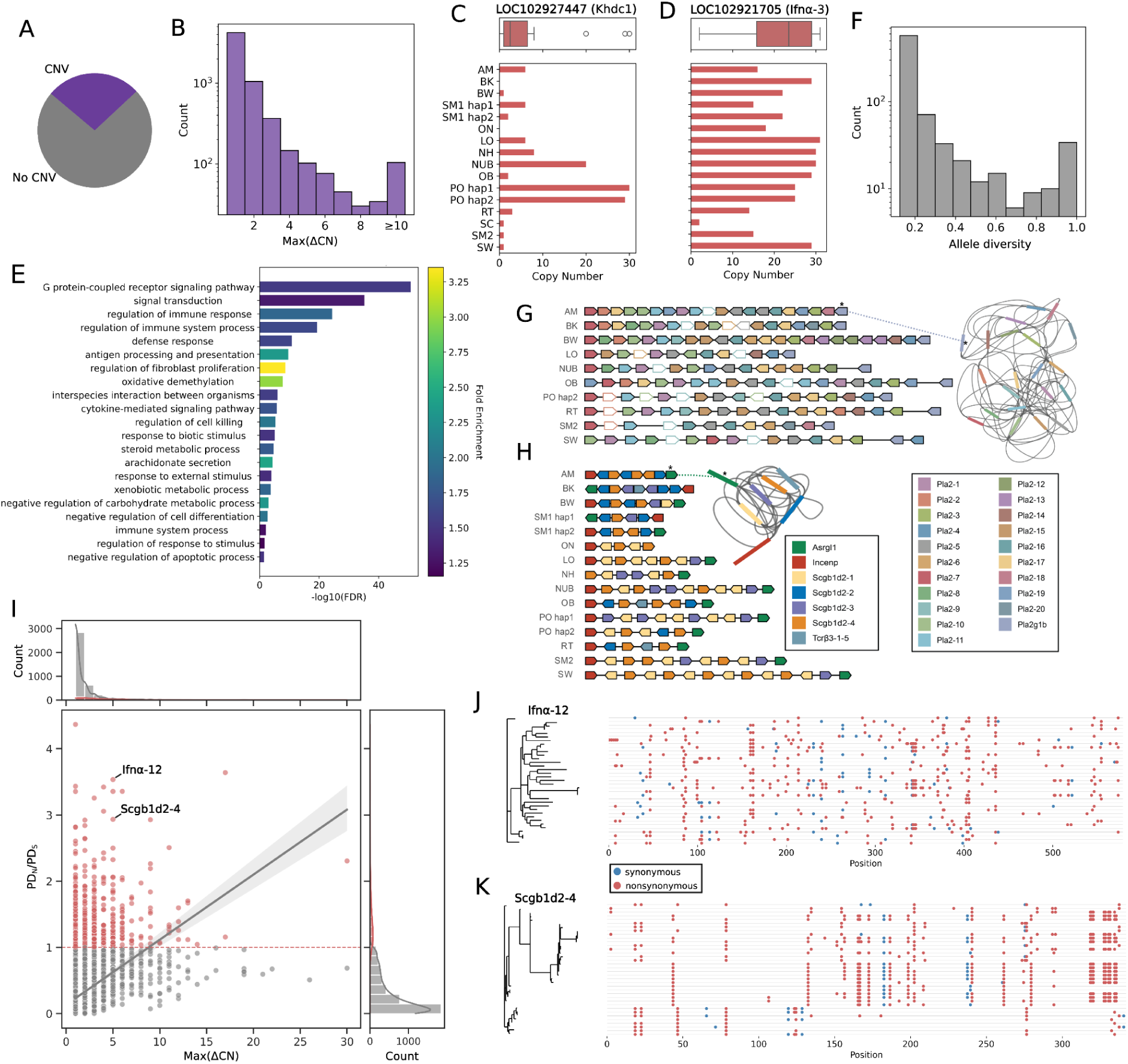
Gene copy number variation and adaptation. (**A**) Proportion of deer mouse genes (N = 6,158/20,956) with copy number variation (purple). (**B**) Distribution of maximum change in copy number across deer mouse haplotypes (Max(ΔCN)) for each gene. **(C-D**) Copy number for (**C)** *Khdc1* and (**D)** *Ifn*⍺*-3* genes among deer mouse haplotypes. Lines inside boxes show the median. Lower and upper bounds for boxes represent the first and third quartile, respectively. Whiskers extend to the minimum and maximum values within 1.5× the interquartile range. (**E**) Representative enriched gene ontology terms for copy number variable genes after redundancy reduction by multidimensional scaling ^55^. (**F**) Distribution of allele diversity, defined as the number of alleles across deer mouse haplotypes normalized by the number of haplotypes with allele calls. (**G-H**) Pangene graphs and linear decomposition of gene order and copy number alleles across deer mouse haplotypes for (**G**) the *Pla2* locus on chromosome 5 and (**H**) the *Scgb1d2* locus on chromosome 1. In (**G-H**), functional copies of genes are filled; pseudogenes are unfilled. For each locus, decomposed haplotypes (left) and corresponding bandage plots (right) are shown. (**I**) Correlation between PD_N_/PD_S_ and Max(ΔCN) for all copy number variable genes. Gray points denote genes with PD_N_/PD_S_ ≤ ∼1 (below red dotted line); red points denote genes with PD_N_/PD_S_ > 1 (above red dotted line). Marginal distributions are shown for each axis. The regression line and 95% confidence interval was inferred using a robust linear model (RLM) with Huber’s T outlier downweighting ^56^. (**J-K**) Maximum likelihood gene tree and multiple sequence alignment for all copies of (**J**) *Ifnα-12* (N = 35) and (**K**) *Scgb1d2-4* (N = 36). Synonymous (blue) and nonsynonymous (red) mutation positions are shown.

Although most genes differ by only one or two copies among haplotypes, copy number variation spans a remarkable dynamic range. More than 100 genes vary by at least 10 copies across deer mouse haplotypes (**Figure 5B**), and many of these expansions or contractions are population-specific. For example, most haplotypes carry relatively few copies of *Khdc1* (median = 2.5), whereas haplotypes from the PO and NUB populations exhibit ten-fold higher copy numbers (**Figure 5C**). Similarly, *Ifnα-3* ranges from only two copies in the SC population to as many as 31 copies in LO, with a median of approximately 24 copies across all haplotypes (**Figure 5D**). These striking, population-specific differences demonstrate that extensive gene copy number evolution is ongoing within deer mice and may contribute to local adaptation.

### Copy number variable genes are important for local adaptation and immune response

Copy number variable genes are strongly enriched for functions involved in environmental sensing and host defense, suggesting that CNVs contribute disproportionately to adaptive variation in natural deer mouse populations. Functional enrichment analyses identified significant overrepresentation of genes involved in sensory perception, environmental interaction, and immune function (**Figure 5E**; Supplementary Table 7). G-protein-coupled receptors were the most enriched category (Q<0.001), including large families of olfactory and taste receptors that are known to evolve through rapid gene gain and loss ^48–51^. We identified 43 olfactory receptor genes that differ by five or more copies among haplotypes, including one canonical olfactory receptor and one vomeronasal receptor whose copy numbers range from 0-12 and 0-13 copies, respectively. Such dramatic variation has the potential to reshape olfactory receptor repertoires and, consequently, which odorants are detected and at what concentrations. Immune-associated pathways, including antigen processing and presentation and cytokine-mediated signaling, were also strongly enriched among CNVs. The largest fold enrichment (>3) was observed for genes regulating fibroblast proliferation, a category composed almost entirely of phospholipase A2 (*Pla*2) genes (Supplementary Table 7). Together, these results suggest that copy number variation preferentially affects gene families involved in environmental sensing and innate immunity.

To examine the haplotype context in which these copy number variants arise, we decomposed pangene graphs into alternative haplotypic paths (“bubbles”). Across 802 loci containing at least one bubble, most exhibited relatively low allelic diversity, with the vast majority (73%) of loci having diversity values below 0.2 (**Figure 5F**; Supplementary Table 8). In contrast, approximately 4.5% of loci exhibited maximal allele diversity, such that every deer mouse haplotype encoded a distinct structural allele. Remarkably, 80% of these highly diverse loci contain immune-related genes ( Supplementary Table 8), consistent with strong selective pressure maintaining structural and haplotypic diversity at genes involved in host-pathogen interactions.

One striking example is the *Pla2* locus on chromosome 5. Although the deer mouse pangenome includes 21 *Pla2* genes at this locus, every haplotype carries a distinct structural allele, resulting in exceptional graph complexity (**Figure 5G**). PLA2 proteins function in antimicrobial defense and inflammatory signaling and have been implicated in resistance to *Borrelia burgdorferi* and Lyme disease pathology ^52,53^. A similarly dynamic pattern occurs at the secretoglobin family 1D member 2 (*Scgb1d2*) locus on chromosome 1, where each deer mouse haplotype contains a unique complement of paralogs (**Figure 5H**). SCGB1D2 has recently emerged as a potent inhibitor of *B. burgdorferi* growth in humans, possibly affecting Lyme disease susceptibility ^54^. Together, these examples illustrate how structural variation reshapes the organization of immune loci in deer mice and identify candidate genes that may contribute to pathogen resistance.

### Molecular signatures of functional diversification

Duplicated genes provide critical substrates for adaptive evolution and functional diversification ^57^. For each copy-number variable gene, we calculated the pairwise differentiation index (PD), the mean number of pairwise differences among all gene copies, corrected for sequence length, and used the ratio of nonsynonymous to synonymous differences (PD_N_/PD_S_) to infer selective pressures acting on duplicated genes ^58^. Approximately 90% of copy-number variable genes exhibit PD_N_/PD_S_ ≤1 (**Figure 5I)**, consistent with purifying selection or neutral evolution ^59,60^. Across genes, greater variation in copy number is associated with higher PD_N_/PD_S_ (RLM, P<0.001; **Figure 5I**), indicating that extensive copy number expansion is generally associated with reduced purifying selection.

A smaller subset of genes (∼10%) exhibits PD_N_/PD_S_ >1, consistent with pervasive positive selection and/or functional diversification. These genes are significantly enriched for immune response regulation and MHC binding (Fisher’s exact test, Q<0.05), and many – including *Ifnα, Pla2* and *Scgb1d2* – which exhibit the highest PD_N_/PD_S_ values also show extensive copy number variation and structural polymorphisms. Pangenomic gene trees and alignments illustrate distinct modes of diversification within this class. For example, *Ifnα-12* (PD_N_/PD_S_ = 3.53; Max(ΔCN) = 5) exhibits numerous nonsynonymous substitutions and long branches separating most gene copies, consistent with ongoing diversifying selection (**Figure 5J)**. In contrast, *Scgb1d2-4* has similarly elevated PD_N_/PD_S_ and Max(ΔCN) values but forms several closely related subfamilies separated by longer branches, suggesting diversification following duplication (**Figure 5K)**. Together, these results indicate that although most duplicated genes remain functionally constrained, a subset of structurally dynamic immune genes accumulates nonsynonymous variation among copies, consistent with functional divergence following duplication.

## Discussion

Assembly-based pangenomes have transformed our understanding of structural variation in humans, livestock, and agricultural crops, yet they have rarely been applied to genetically and ecologically diverse populations from the wild. Here, we present the pangenome for the deer mouse, providing a genome-scale view of structural variation across natural populations of a wild mammal. More than one billion base pairs of sequence present in deer mouse haplotypes are absent from the traditional deer mouse reference genome (BW), demonstrating that a single reference captures only a fraction of the genomic diversity within this species. This missing sequence has important implications for association mapping, population genetic inference, and functional genomics studies employed in the deer mouse model ^28,61–68^. Moreover, replacing one reference genome with another does not eliminate this bias: approximately 60% of the pangenomic sequence is accessory sequence across the 16 deer mouse haplotypes analyzed here. Although estimates of accessory sequence are influenced by differences in sample, genome contiguity, and pangenome construction methods, this proportion is substantially higher than reported for other mammals, such as 28% across 16 human haplotypes ^11^, ∼5% across 15 sheep breeds ^14^, and 2.8% for 6 cattle genomes ^15^. Together, these observations suggest that deer mice harbor exceptional structural genomic diversity. Whether this degree of diversity is unusual or representative of wild mammals more broadly remains an open question that will require pangenomes from additional natural populations of mammals.

The majority of structurally variable sequence in the deer mouse pangenome is repetitive and reflects ongoing expansion of endogenous retroviral elements. More strikingly, however, structural variation extends well beyond repetitive DNA to encompass nearly one-third of annotated protein-coding genes. We find that 29% of genes annotated in the canonical deer mouse reference exhibit copy-number variation across deer mouse haplotypes, with some loci displaying levels of copy number and gene order variation unprecedented among previously studied mammals ^47,69,70^. This previously cryptic variation has been largely inaccessible to evolutionary genetics and functional genomic studies despite growing evidence that CNVs frequently contribute to local adaptation, phenotypic diversification and host-pathogen interactions. The enrichment of copy number variation among genes involved in sensory perception, environmental interactions, and immunity suggests that structural variation has been an important substrate for adaptation during the remarkable ecological diversification of deer mice across North America ^71^.

This same genomic flexibility may also contribute to the species’ ability to serve as a reservoir for numerous zoonotic pathogens, leading to disease, including hantavirus pulmonary syndrome, Lyme disease, and viral encephalitis ^35^. A particularly intriguing finding is the positive association between copy number variation and PD_N_/PD_S_. Although most duplicated genes remain under strong purifying selection, a subset of structurally dynamic immune genes exhibit signatures consistent with positive selection at both the structural and molecular levels. These observations suggest that copy number expansion and sequence diversification may proceed in concert at genes involved in host-pathogen interactions, although determining the evolutionary mechanisms responsible for these patterns will require future population genetic and functional studies.

Beyond providing an important resource for the deer mouse community, this work demonstrates that natural populations can harbor substantially greater structural genomic diversity than is apparent from a single mammalian reference genome. Our results reveal remarkable plasticity in mammalian gene repertoires and suggest that structural variation is concentrated in gene families mediating interactions between organisms and their environments. More broadly, these findings illustrate the power of pangenome approaches to uncover previously hidden substrates for adaptation and emphasize the importance of extending pangenome analyses to genetically and ecologically diverse species in the wild.^25^.

## Methods

### Sampling and sequencing

To maximize genetic, geographical, and ecological variation, we selected deer mice from different habitats across their range (Figure 1A; Table S1). We previously generated several assemblies for diverse deer mouse populations (BW, NUB, SW, BK). We selected 9 additional populations for *de novo* assembly. These include several populations that had been derived from the wild to establish breeding colonies in the Hoekstra lab: *P. m. sonoriensis* (SM2) from the foothills of the Sierra Nevada, *P. m. rufinus* (RT) from the Monzano Mountains of New Mexico, the forest-dwelling *P. m. artemisiae* (AM) and prairie-dwelling *P. m. osgoodi* (OB) from western Montana. Since *Peromyscus polionotus* is phylogenetically nested within *Peromyscus maniculatus*, we also incorporated *P. polionotus leucocephalus* (LO) from the Florida panhandle and *P. polionotus subgriseus* (PO) from central Florida. In addition, we also obtained museum specimens or previously collected nonlethal tail clips for *P. m. gracilis* from the forests of southern Ontario (ON), *P. m. bairdii* from New Hampshire (NH), and *P. m. santacruzae* from Santa Cruz Island, California (SC) (**Figure 1B**; Supplementary Table 1). We also sequenced a single individual from an outgroup species, *P. leucopus*, sampled from Black Mountain, Kentucky.

All assemblies were produced using PacBio HiFi sequencing, with coverage ranging from 13X to 33X, except for SC, which was produced using Oxford Nanopore Technologies (30X coverage) due to limited input material (Supplementary Table 2). For the AM, SM2, OB, and LO populations, we used a previously described approach ^34^ in which we applied PacBio HiFi and Omni-C proximity ligation sequencing to inter-population F1 hybrids (AMxSM2, OBxLO). After scaffolding, we used ancestry-informative SNPs to assign chromosomes to each source population (see details below; Extended Data Figure 1). For the remaining populations, we sequenced a single individual from each population. Fresh tissue was retrieved from each mouse immediately after euthanization for SM2, RT, OBxLO, AMxSM2 and PO –derived from laboratory colonies maintained at Harvard University. The SC, NC, and ON tissue samples were collected from wild-caught individuals and preserved in ethanol.

### Long-read sequencing

We extracted high molecular weight DNA from each sample using the Pacific Biosciences Nanobind kit (PacBio, Menlo Park, CA, USA) following the manufacturer’s protocol. We quantified DNA samples using a Qubit 2.0 Fluorometer (Life Technologies, Carlsbad, CA, USA) and Femto Pulse (Agilent, Santa Clara, CA, USA) and constructed PacBio SMRTbell libraries (∼20 kb) using a SMRTbell Express Template Prep Kit 2.0 (PacBio, Menlo Park, CA, USA). We performed Pacbio sequencing using a combination of Sequel II and Revio systems. For Sequel II, we used the Sequel II Binding Kit 2.0 (PacBio, Menlo Park, CA, USA) to bind the library to polymerase. In the case of Revio, we used the Revio polymerase kit (PacBio, Menlo Park, CA, USA). For the degraded SC sample, we performed size selection to eliminate small DNA fragments prior to nanopore sequencing using a Circulomics SRE kit (Circulomics, Menlo Park, CA, USA), which selects for fragment sizes ≥10 kb. Then, we generated a nanopore sequencing library using the SQK-LSK109 kit (Oxford Nanopore Technologies) and loaded it onto a PromethION (Oxford Nanopore Technologies) for sequencing. All sequencing was conducted at the Bauer Core Facility at Harvard University.

### Omni-C sequencing

For PO, AMxSM2, and OBxLO samples Omni-C libraries were prepared from flash-frozen liver, brain, and muscle tissues following the Dovetail Genomics mammalian tissue protocol (see https://dovetailgenomics.com/wp-content/uploads/2021/09/Omni-C-Protocol_Mammals_v1.4.pdf). Chromatin was fixed *in situ* using formaldehyde and digested with DNase I. The resulting chromatin ends were repaired and ligated to a biotinylated bridge adapter, followed by proximity ligation. After reversing crosslinks and purifying the DNA, non-internal biotin was removed. Sequencing libraries were generated using NEBNext Ultra enzymes and Illumina-compatible adapters. Biotinylated fragments were enriched using streptavidin beads, PCR-amplified, and sequenced to approximately 30x coverage on an Illumina Novaseq platform.

### Iso-Seq for genome annotation

We collected six tissues from a single BW individual and pooled them for an Iso-Seq experiment: liver, brain, testes, spinal cord, vomeronasal organ (VNO) and main olfactory epithelium (MOE). Liver, brain and testis tissues were equally represented in the Iso-Seq data. Spinal cord was represented at 1/2 the coverage of liver, and VNO & MOE each at 1/4 the coverage of liver. Total RNA was extracted from fresh frozen tissue samples using Qiagen RNeasy Plus Universal mini kit (Qiagen, Hilden, Germany). RNA samples were quantified using Qubit 2.0 Fluorometer (Life Technologies, Carlsbad, CA, USA), and RNA integrity was checked with TapeStation 4200 (Agilent Technologies, Palo Alto, CA, USA). For Iso-Seq library construction, 300 ng of total RNA was converted into amplified full-length cDNA using the SMARTer PCR cDNA synthesis Kit (Clontech, Mountain View, CA, USA). The library for PacBio Sequel was constructed using SMRTbell Express Template Prep Kit 3.0 (PacBio, Menlo Park, CA, USA). The library was bound to polymerase using the Sequel Binding Kit (PacBio) and loaded onto PacBio Sequel using the MagBead Kit V2 (PacBio). Sequencing was performed on 1 PacBio Sequel II SMRT Cell (24-hour movie time). After sequencing, CCS reads were obtained using the CCS algorithm within PacBio ccs (version 4.2.0) ^72^, run using the default parameters. We used Lima (version 2.9.0) to remove cDNA primers (https://github.com/PacificBiosciences/barcoding), followed by iso-seq refine (version 4.0.0) (https://isoseq.how/) to remove polyA tails and artificial concatemers. Then, we used isoseq cluster (version 4.0.0) (https://isoseq.how/) to cluster *de novo* isoforms. The BW reference genome assembly was annotated by NCBI using the Iso-Seq data produced in this study.

### Genome assembly and quality control

All samples sequenced with PacBio technologies were assembled using hifi-asm (version 0.19.8) ^73^ with default parameters, but setting the expected haploid genome size (--hg-size) to 2.6 Gb and lowering the similarity threshold for duplicate haplotigs (-s) at the read level to 0.4, based on improved assembly statistics after optimizing these parameters for our dataset. We also included Omni-C data in our hifi-asm command for PO, AMxSM2, and OBxLO to produce phased diploid assemblies. To scaffold AM, SM2, OB, and LO assemblies, we first aligned Omni-C data to each assembly using bwa mem (version 0.7.18) ^74^ with the flag-5SP. Then, we marked duplicate and split reads using samblaster (version 0.1.26) ^75^ followed by samtools view (version 1.11) ^76^ with flags-@ 14-S-h-b-F 3340. We filtered the subsequent bam files using the HapHiC utility tool filter_bam (version 1.0.6) ^39^ with flags 1 --nm 3 and scaffolded assemblies using HapHiC (version 1.0.6) ^39^ with default parameters and an expected diploid chromosome number of 48. Then we evaluated ancestry across the phased genomes using ancestry-informative sites. Specifically, we identified divergent SNPs (Fst > 0.9 for LO v. OB, Fst > 0.75 for AM v. SM2) using whole-genome short-read sequencing from AM, SM2, OB, and LO populations. We then mapped the AM, SM2, OB, and LO genome assemblies to the Pman2.1.3 reference genome (GCA_003704035.3) using minimap2 (version 2.9) ^77^ and extracted each assembly’s allele at ancestry-informative SNPs using samtools (version 1.10) mpileup ^76^. We computed population allele frequencies in 500 kb windows using the alleles present in the genome assembly of interest to visualize average ancestry across each assembly. PacBio-based primary assemblies were screened for adaptor and contaminant sequence with NCBI FCS-adaptor and FCS-GX (https://github.com/ncbi/fcs). For SC, which was sequenced with Oxford Nanopore Technology, we removed adaptors using porechop_abi (version 0.5.0) ^78^ with the parameter --ab_initio and performed genome assembly using Flye (version 2.9.2) ^79^ with parameters --nano-hq --genome-size 2.6g --scaffold. We also used Flye to polish the resulting SC assembly with the --polish-target parameter. We computed assembly quality metrics using gfastats (version 1.3.6) ^80^, QUAST (version 5.2.0) ^81^ with parameter --split-scaffolds, and compleasm (version 0.2.9) ^82^ with parameter-l mammalia --odb odb10 (mammalia_odb10, release 2024-01-08). Gfastats and QUAST provide a range of contiguity metrics such as N50 and auN, whereas compleasm uses BUSCO gene sets to assess assembly completeness. Assembly statistics for all assemblies are reported in Supplementary Table 2. Although we produced phased and primary assemblies for all PacBio samples, we found that phased assemblies consistently displayed lower contiguity and completeness than primary assemblies. In addition, we observed high levels of duplicated genes in our initial ON and NH assemblies. Thus, we performed a post hoc purge duplication step on these two assemblies with purge_dups (version 1.2.6) (https://github.com/dfguan/purge_dups), which drastically improved the BUSCO-based assessment of gene set completeness/duplication levels. We required a minimum mammalian BUSCO completeness score of 89% for assembly consideration. Only phased assemblies for F1 individuals (AMSM2 and OBLO), PO, and *P. leucopus* passed this threshold. As a result, we used primary assemblies for NH, OT, RT, SM2, and SC.

### Repeat annotation

To annotate repeats in each genome assembly, we first retrieved lineage-specific curated TE models and satellite sequences previously mined for *P. maniculatus* ^32^. We combined this lineage-specific library with all curated ancestral *Peromyscus* TE models available from DFAM ^83^. Then, we ran RepeatMasker (version 4.1.6) ^84^ on each assembly with parameters-e ncbi-excln-s-no_is-u-noisy-html-xm-a-xsmall-source using our combined TE library. We resolved overlapping or redundant repeat annotations based on highest alignment scores using the RepeatMasker utility script RM2Bed.py with the parameter --ovlp_resolution highest_score^84^.

### Sequence partitioning

Since PGGB performs all-vs-all comparisons to generate unbiased pangenome graphs, it is generally not feasible to run PGGB on all sequences for a large, complex genome ^10^. Instead, we partitioned sequences into chromosomes and ran PGGB separately for each chromosome ^10,11^. To do this, we first gathered all contigs from all selected assemblies into one fasta file using the PanSN-spec naming pattern (https://github.com/pangenome/PanSN-spec). Then, we ran wfmash (version 0.13.0) ^85^ with parameters-p 85-n 18 to generate all-vs-all alignments across all contigs. Since deer mouse centromeric satellites share high levels of structural and sequence similarity across chromosomes and populations ^34,44^, we masked centromeric sequences from these alignments. Then, we projected alignments into graph networks using the PGGB utility script paf2net.py. Finally, we implemented the Leiden algorithm ^86^ through the PGGB utility script net2communities.py to identify communities from our alignments. We visualized these results using igraph ^87^ through python, finding 24 large communities corresponding to individual chromosomes as annotated in the chromosome-level assemblies for BW, NUB, BK, and SW. Smaller communities or singletons that did not clearly cluster with any chromosome were ∼85% repeats on average, nearly twice as repetitive as contigs that clustered with chromosomes (∼45%); we removed these contigs from downstream analyses.

### PGGB pangenome graph construction

We used PanGenome Graph Builder (PGGB) to construct unbiased pangenome graphs for each deer mouse chromosome ^10^. PGGB is one of two currently available pipelines for nucleotide-level pangenome graph construction that scales to large, complex eukaryotic genomes. The other is minigraph-cactus ^9^. Both produce similar results, as demonstrated by the Human Pangenome Reference Consortium ^11^. However, unlike PGGB, minigraph-cactus fails to incorporate repetitive regions in the resulting graph. Thus, we opted to use PGGB for constructing the pangenome graph. We ran PGGB on all contigs across all considered genome assemblies separately for each chromosome community, with the parameters-p 85-V BW:#-n 19-k 23-m, per the developers’ recommendations. PGGB produces several final files including a final pangenome graph in odgi format ^42^, a final pangenome graph in GFA format and a VCF containing variant calls generated using VG deconstruct ^88^.

### Minigraph-cactus pangenome graph construction

We also generated a minigraph-cactus (MC) ^9^ pangenome graph since the PGGB graph can be intractable for mapping resequencing data. We used a previously developed snakemake pipeline to run MC (version 2.9.9) on the Harvard FASRC cluster https://informatics.fas.harvard.edu/resources/tutorials/pangenome-cactus-minigraph/ with default parameters and BW as the backbone reference assembly.

### Pangenome evaluation

We used odgi squeeze to concatenate PGGB graphs from each chromosome into one final pangenome graph. Then, we ran odgi stats ^42^ to obtain graph metrics, such as the number of nodes and the overall length of the graph, for both the PGGB and MC graphs. For the PGGB graph, to assess the extent of missing sequence in the canonical deer mouse reference (BW), we compared the total length of our pangenome graph in nucleotide space with the length of the sequence represented by BW. We then recomputed this metric excluding outgroups by extracting a graph that only represented *P. maniculatus* haplotypes using odgi extract ^42^ with options-p and-b to specify genomic paths and regions corresponding to deer mouse haplotypes. We estimated the average proportion of pangenomic sequence contributed by individual deer mouse haplotypes using odgi heaps ^42^ with parameters-H-d 1-n 100. We identified private sequences, i.e., sequences represented in the graph by only one haplotype, using gretl core ^89^ after converting our final concatenated pangenome graph into GFA format with odgi view ^42^. To evaluate pangenome growth statistics, we ran panacus histgrowth (version 0.2.3) ^90^ on the final concatenated GFA graph with parameters-l 1,2,1-q 0,0,1-t10-H-c all-a-o html. We ran the same command to assess growth of the MC pangenome. We extracted core and accessory sequences from the pggb graph in fasta format using gfatools gfa2fa (version 0.5-r292) (https://github.com/lh3/gfatools). Then, to evaluate repeat composition in the core and accessory genome, we ran RepeatMasker with the same parameters mentioned previously.

### Tree construction and comparison

To reconstruct the evolutionary relationships among haplotypes based on small variants, we first used bcftools to filter our VCF file for biallelic SNPs with no missing data (view --min-ac 1:minor --type snps --max-alleles 2-i’F_MISSING==0.0) and then further thinned the callset for one maker every 10 kb (+prune-w 10000bp-n 1-N rand) to reduce the effect of linkage. We then used three different approaches to build phylogenetic trees. First, we used tetrad, a quartet-based approach implemented in ipyrad (version 0.9.105) ^91^, evaluating all possible quartets and statistical support with 100 bootstrap replicates. Second, we used custom python code and the DistanceMatrix and nj functions in the skikit-bio package (version 0.7.0) ^92^ to build a Neighbor-Joining tree, randomly resampling sites 100 times for bootstrap evaluation. Finally, we used the concatenated SNP matrix in IQ-Tree (version 3.0.1) ^93^ to generate a Maximum-Likelihood tree specifying 1000 ultrafast bootstraps with the nearest-neighbor interchange optimization step (-B 1000 --bnni) to evaluate support.

To build trees based on the variant graph structure, we calculated the Jaccard distance between paths in the PGGB and MC graphs using odgi similarity with flags --delim=# --delim-pos=1. Then, we converted Jaccard distance matrices to newick trees using python. We used R to compare and plot SNP-based, PGGB and MC trees. We computed Robinson-Foulds distance and plotted co-phylogenetic plots using phytools through R ^94^. We used a custom approach in R to compare the position of each haplotype between trees. First, we rooted each tree at *P. eremicus* and standardized branch lengths by assigning each branch length a value of 1. Then, we computed the positional depth *d* for each leaf in each tree, such that *d* = 5 means the species is 5 splits away from the root. Finally, we computed leaf displacement Δ*d* for each leaf wherein Δ*d* = |*d*_tree1_- *d*_tree2_|. We used Kendal’s test to test for a correlation between private sequence and leaf displacement between PGGB and SNP-based trees across deer mouse haplotypes.

### Pangenome graph visualization

We generated one-dimensional and two-dimensional visualizations for each PGGB chromosome graph, as shown in Figure 2A and 2B, respectively. To produce two-dimensional visualizations, we ran odgi draw (version 0.8.6) ^42^ with parameters-H 1000 using layouts produced from PGGB. We produced compressed one-dimensional visualizations displaying depth across all paths along the graph for each chromosome using odgi viz ^42^ with parameters-x 1500-y 500-a 10-O-I Consensus.

### Variant filtering and processing

PGGB directly calls single nucleotide polymorphisms, indels (variants ≤50 bp), and unbalanced structural variants (variants >50 bp) from pangenome graphs using VG deconstruct ^88^. However, these variant calls require further processing for downstream analyses ^11^, which we did following recommendations from the Human Pangenome Reference Consortium ^11^. Specifically, we first indexed VCFs produced with VG deconstruct for each chromosome using tabix (version 1.11) ^95^. Next, we removed large spurious variants (>100000 bp) using vcfbub (version 0.1.0) (https://github.com/pangenome/vcfbub) with parameters-l 0-r 100000. Then, we ran vcfwave ^96^ with parameter-I 1000 to decompose complex overlapping alleles into simpler primitive variants, followed by bcftools norm-m+both (version 1.9) ^97^ to join biallelic variants into multiallelic variants separately for SNPs and indels. We concatenated VCF files from each chromosome using bcftools concat ^97^ and sorted it using bcftools sort ^97^. Finally, we removed variant calls in regions of the graph with low and high node depth. To do this, we calculated node depth in the PGGB graph across 1 kb windows using odgi depth ^42^. Then, we filtered all variant calls in regions with node depth <2 or ≥1000 using bedtools intersect (version 2.29.1) ^98^ with options-header-v.

### Variant polarization

We separated biallelic and multiallelic sites in our variant calls using bcftools view ^97^ with options-m2-M2 since we focused primarily on biallelic sites in several downstream analyses. To polarize variants, we first filtered our VCF for outgroup genotypes using bcftools view with the-s parameter ^97^ and extracted genotypes at each site in tab delimited format using bcftools query ^97^. Then we polarized allele calls based on genotypes in outgroup samples. Specifically, for a site to be considered for polarization, we required that either three of the four outgroup genomes possess the same genotype or two of the four outgroup genomes possess the same genotype in cases where the remaining two outgroup genomes possessed missing genotypes. We then annotated the ancestral alleles for all sites that fulfilled these requirements and for which genotyping information was available. We marked polarization results for all other sites as ambiguous. We added ancestral allele information to our VCF using bcftools annotate ^97^ and generated ingroup VCFs excluding genotypes for outgroup samples using bcftools view ^97^.

### Variant classification

We classified variants based on the size of reference and alternate alleles. We note that VG deconstruct does not currently report balanced structural variants such as inversions ^16,88^. Thus, we limited our analysis to single nucleotide polymorphisms, multinucleotide polymorphisms (MNPs), indels and unbalanced structural variants. Variants for which both the reference and alternate allele were 1 bp were classified as SNPs. If the reference and alternate allele had equal sizes <50 bp, we classified them as MNPs. For polarized variants in which the reference and alternate alleles differed in size, if the ancestral allele was larger than the derived allele, we classified the variant as a deletion. Otherwise, we classified the variant as an insertion. Insertions and deletions were further classified as either indels or SVs based on their size: if the maximum size of all alleles for a given site was ≤50 bp, we classified the variant as an indel; if the maximum size was >50 bp, we classified the variant as a structural variant. Finally, we classified indel insertions, indel deletions, SV insertions, and SV deletions as complex when both ancestral and derived alleles were longer than 1 bp.

### SV repeat composition and TE insertion analysis

To identify the repeat composition of SVs, we first focused on all SVs including multiallelic SVs. For multiallelic SVs, we only considered the longest allele to minimize redundancy in our repeat composition analysis. Then we ran RepeatMasker with the same parameters and repeat library as for assembly-wide repeat annotation. We also did this separately for all polarized SV insertions and deletions to estimate the relative repeat composition of insertions and deletions. To identify potential polymorphic TE positions, we filtered for SV insertions and deletions that were 60% covered by models for a particular TE family.

### Analysis of high-depth regions

We used odgi depth with default parameters to identify 100 kb regions of the PGGB graph that had high node depth, projected onto the BW genome assembly. For this analysis we considered high-depth regions as any 100 kb region with node depth >500. We sorted and merged high-depth regions within 100 kb of each other using bedtools. Then, we performed permutation tests for enrichment of different repeats in high-depth regions using GAT (version 1.3.5) ^99^ with parameters --nbuckets=10 --bucket-size=100000. We visualized depth distributions across chromosomes with Rideogram ^100^.

### Analysis of PMsat positional variation and chromosomal rearrangements

To compare PMsat array positions across chromosomes for all deer mouse haplotypes, we first used ragtag to scaffold genome assemblies not scaffolded with Omni-C data. These samples included SM1 hap1, SM1 hap2, PO hap1, PO hap2, SC, ON, RT, and NH. We used ragtag (version 2.1.0) ^101^ with parameters --mm2-params’-x asm20’ to scaffold these assemblies with the BW assembly as the reference. We removed unplaced scaffolds from each assembly using bioawk (version 20110810) (https://github.com/lh3/bioawk) and mapped to the BW assembly using minimap2 (version 2.30-r1287) ^77^ with flags-ax asm20 --eqx. Then, we used paftools.js sam2paf (version 2.30-r1287) ^77^ to convert sam files to paf files, and generated dotplots to visualize synteny between each assembly and the BW reference assembly. We manually inspected dotplots to identify and correct misoriented scaffolds. We mapped PMsat ^44^ to each correctly scaffolded assembly using RepeatMasker with the same parameters as previously. We merged PMsat arrays within 100 kb of each other using bedtools merge. Then, we visualized variation in PMsat positions across chromosomes and haplotypes by normalizing PMsat coordinates by chromosome length for each haplotype. We were also interested in PMsat array positions relative to large chromosomal rearrangements, since rearrangements might play a role in mediating PMsat repositioning ^34^. Due to the previously noted challenges of extracting large chromosomal rearrangements such as chromosomal inversions from pangenome graphs ^102^, we instead used SyRi ^103^ to identify chromosomal rearrangements between haplotypes. We focused this analysis on assemblies scaffolded with Omni-C data (BW, BK, SW, NUB, AM, SM2, OB, and LO) and ignored ragtagged assemblies because inversion breakpoints often occur at contig breaks ^34^. Thus, ragtagged assemblies likely fail to capture many true inversions present in respective haplotypes. We ran Syri (version 1.7.1) with parameters-k-F S and visualized chromosomal rearrangements and PMsat positions across haplotypes using plotsr (version 1.1.1) ^104^ with flags --itx --log DEBUG-d 600-S 0.5-W 8-H 8-f 8.

### Gene copy number and order variation

Although PGGB is an excellent tool for unbiased pangenome graph construction at nucleotide-level resolution, the complexity of PGGB graphs makes it challenging to study gene copy number and order variation in a high-throughput manner. This can be particularly problematic for paralogs, which are sometimes collapsed in PGGB graphs, or structural variants affecting genes in repeat-rich genomic regions due to the complex bubbles formed by multiple-sequence alignments of repeats. The recently developed tool, pangene ^46^ overcomes these challenges by constructing a bidirected gene graph, in which each node is a gene and each edge encodes the adjacency and orientation of that gene relative to its neighbor. To implement pangene, we first generated gene annotations for each deer mouse haplotype using miniprot (version 0.12-r237) ^105^ with options --outs=0.95 --no-cs-Iut16 and all deer mouse proteins as input. Then, we ran pangene (version 1.1-r231) ^46^ with options-c 100 to generate a pangene graph in GFA format. We also reran pangene with the additional option-bed=flag to produce a bedfile of gene annotations along each contig in our pangenome, as well as useful information pertaining to each annotation, such as the presence of premature stop codons and divergence estimates relative to the source protein. We used the pangene utility script pangene.js gfa2matrix to identify CNVs between haplotypes and removed genes on the X chromosome. We ran pangene.js call ^46^ to identify bubbles in the pangene graph in homologous regions across haplotypes. This analysis was limited to genes and haplotypes harbored within a single contig and alleles fragmented across multiple contigs for a given haplotype were discarded. We parsed pangene bubbles to obtain allele counts and composition, and discarded loci with multiple nested bubbles in which the same haplotype carried multiple alleles. We then manually visualized allelic variation for gene copy number and order using pangene’s gfa-server ^46^ as well as bandage (version 0.8.1) ^106^.

### Gene ontology enrichment analysis

We performed gene ontology (GO) enrichment analyses for copy number variable genes using goatools ^107^ through python and corrected for multiple hypothesis testing using the Benjamini/Hochberg method ^108^. We reduced redundancy in the resulting set of enriched GO terms using Revigo (version 1.8.1) ^55^, clustering terms with dispensability values >0.7 and prioritizing terms with lower Q-values.

### Molecular evolution analysis

To test for patterns of molecular evolution in copy number variable genes, we first generated multiple sequence alignments from all copies for each copy number variable gene using MAFFT (version 7.50) ^109^. Then, we calculated the pairwise differentiation index (PD) ^58^, the mean number of pairwise differences for all copies of a gene corrected for the number of sites, for all copy-number variable genes across deer mouse populations. We did this separately for nonsynonymous (PD_N_) and synonymous (PD_S_) polymorphisms. We then calculated PD_N_/PD_S_ for each gene with PD_S_ >0.01. We interpreted values of PD_N_/PD_S_ ≤1 as likely evidence of negative selection or neutrality and PD_N_/PD_S_ >1 values as evidence of diversifying or balancing selection ^58,110,111^. Robust linear regressions (RLMs) associating PD_N_/PD_S_ and variation in copy number across genes were performed using python with Huber’s T outlier downweighting to account for outlier effects ^56^.

### Obtaining biogeographical data for Figure 1

We obtained geographical boundaries for all level I terrestrial ecoregions (the broadest level of ecoregion grouping) from the North American Environmental ATLAS (http://www.cec.org/; last accessed August 29, 2024).

## Data availability

We retrieved previously published high-quality deer mouse genome assemblies, as well as outgroup species genomes from NCBI. These include chromosome-level assemblies produced from F1 hybrids for BW, NUB, BK, and SW ^34^ (NCBI: GCA_049852395.1, GCA_049852375.1, GCA_049852355.1, and GCA_049852335.1, respectively), phased assemblies produced as part of the California Conservation Genomics Project for SM1 (NCBI: GCA_026167925.1 and GCA_026229955.1) ^112^, and publicly available primary assemblies for *Peromyscus leucopus* (GCF_004664715.2) ^113^ and *Peromyscus eremicus* (GCF_949786415.1). We retrieved published data to produce pangenome growth curves for scrub jay ^26^, humans ^26^ and cichlid fish ^114^. Raw data generated in this study is publicly available from NCBI’s Sequence Read Archive (BioProject: PRJNA1531221). Genome assemblies generated in this study will be deposited to NCBI before publication.

## Code availability

No new software packages or algorithms were developed for this research.

## Acknowledgments

Consistent with Nature’s policy, AI-assisted tools were used to suggest refinements to the language of the manuscript. The authors remain fully accountable for its intellectual content. The authors thank John Orrock, Jim Kruppa, Rebecca Rowe and Joshua Willems, as well as Virginie Millien, Kirsten Crandall, and Anthony Howell (Redpath Museum) for help with collecting or kindly sharing tissue samples. We thank Erik Garrison and Andrea Guarracino for their insights into optimal parameters for running PGGB on our dataset. We thank Thomas Keane for permission to use the *P. eremicus* assembly in our analysis. We also thank Bohao Feng, Tim Sackton, Russell Corbett-Detig, Scott Edwards, Shuonan He, and Runyang Nicolas Lou for helpful discussions as well as feedback on this manuscript.

## Author contributions

L.G., A.F.K., and H.E.H conceived the study and selected samples for sequencing. C.K. and A.F.K. performed sample preparation. L.G, O.H., and A.F.K. performed genome assembly and quality control. L.G. generated pangenome graphs and performed all pangenome analyses, whole-genome comparison, and CNV analyses with input from A.F.K. and P.S. L.G., A.F.K., and P.S. wrote the original draft of the manuscript. All authors read and approved the final manuscript.

