## Supplemental Figures for "Pangenome analyses uncover the scale and evolutionary consequences of structural variation in a wild mammal"

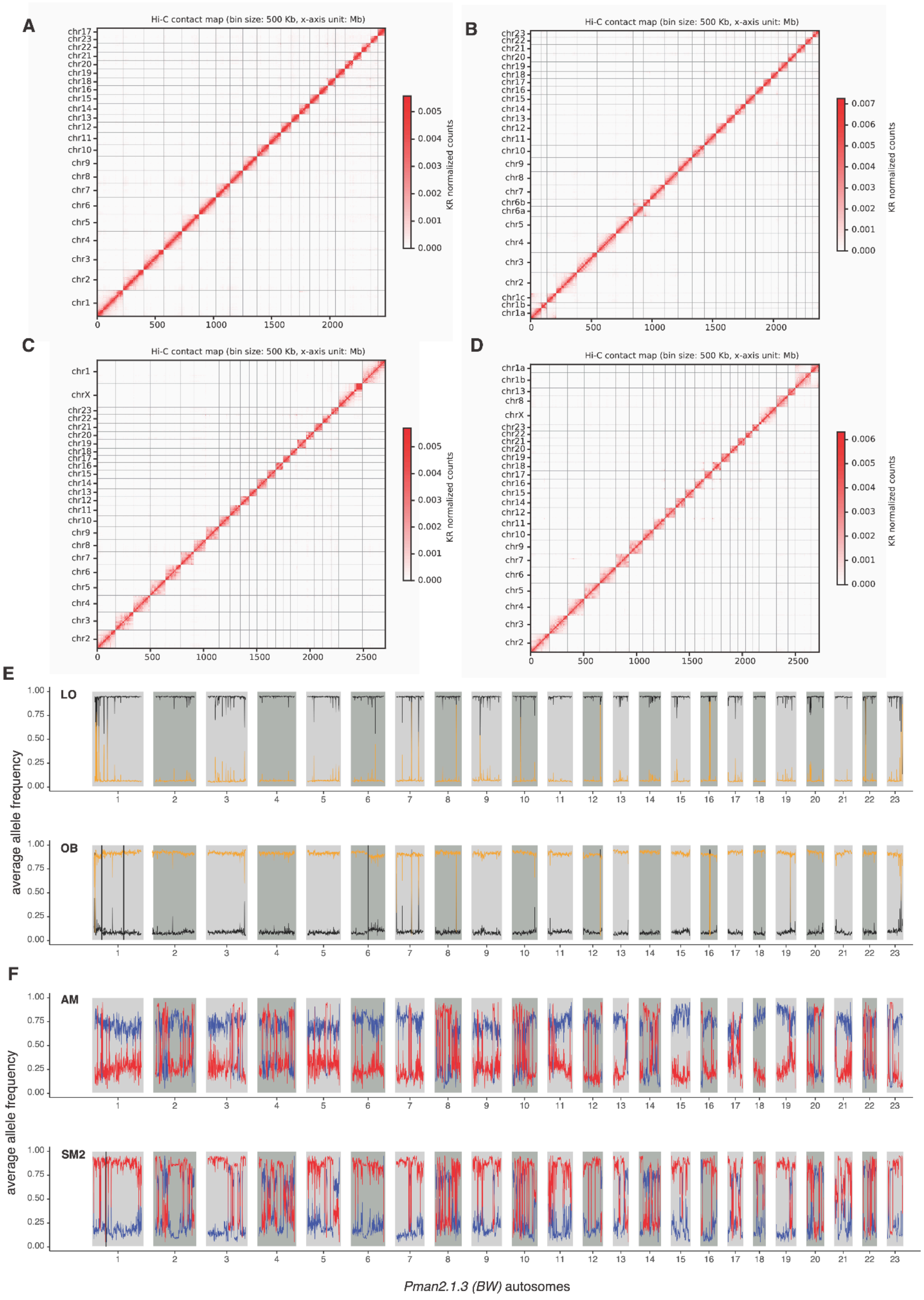


**Extended Data Figure 1:** Contact maps for Omni-C data aligned to assembled (**A**) LO, (**B**) OB, (**C**) AM, and (**D**) SM2 chromosomes. Ancestry plots for **(A)** phased OBxLO and (**B**) AMxSM2 genome assemblies. (**A**) LO and OB ancestry are shown in black and yellow, respectively. (**B**) AM and SM2 ancestry are shown in blue and red, respectively.

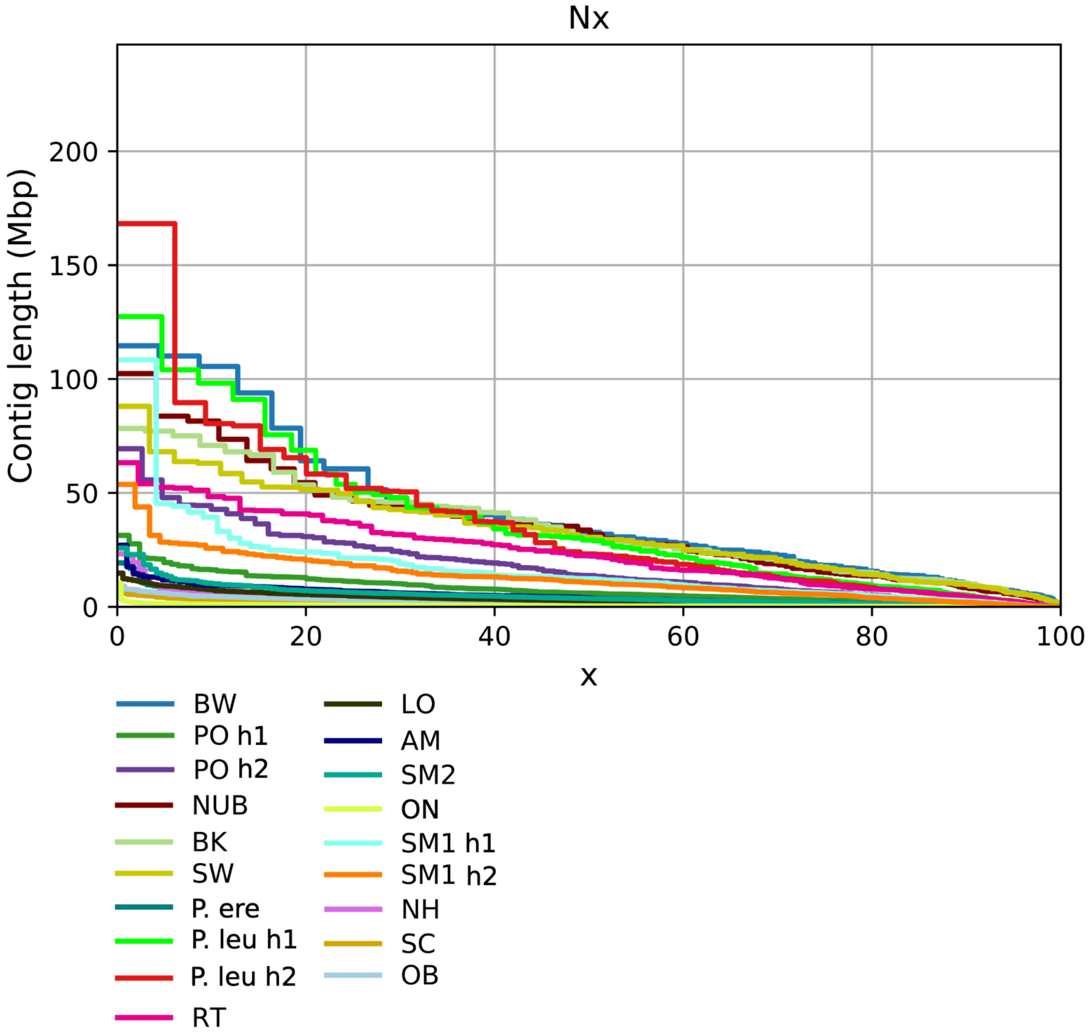


**Extended Data Figure 2:** Nx plot showing the percent of the genome occupied by contigs from largest to smallest for each assembly selected for pangenome construction.

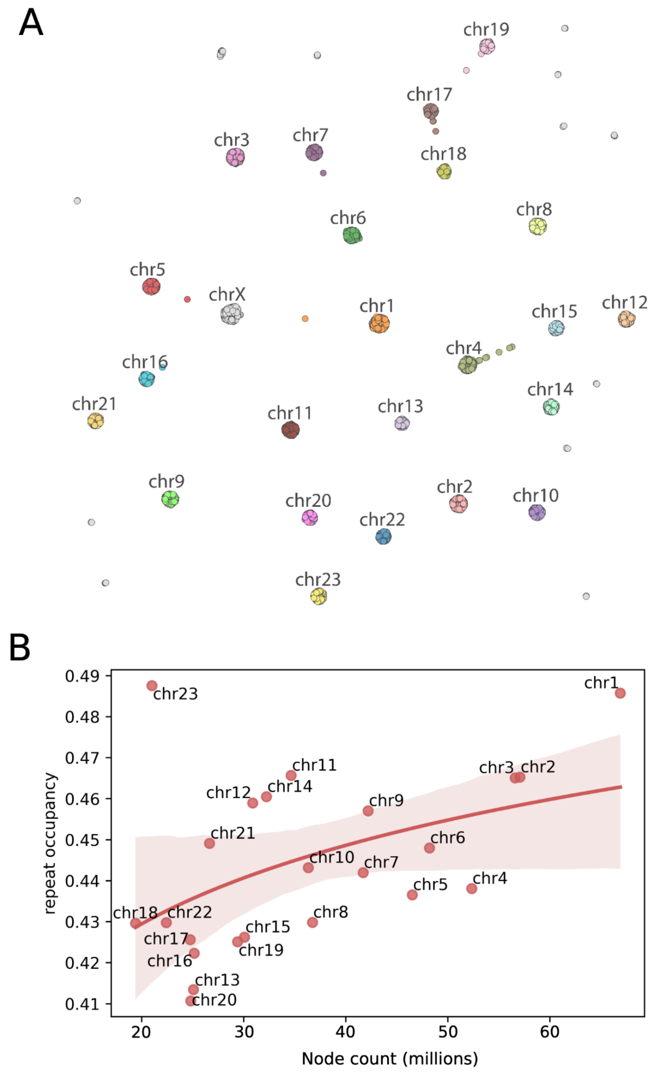


**Extended Data Figure 3:** (**A**) Contig clustering generated from all-vs-all alignments. Each node represents a contig. Most contigs clearly cluster with distinct chromosomes as annotated in the previously-generated chromosome-level assemblies of BW, NUB, BK and SW. (**B**) Correlation between node count and repeat occupancy across deer mouse chromosomes (Kendall's Tau = 0.38, P = 0.01). Each chromosome is labeled as an independent point as well as a linear regression line and 95% confidence intervals.

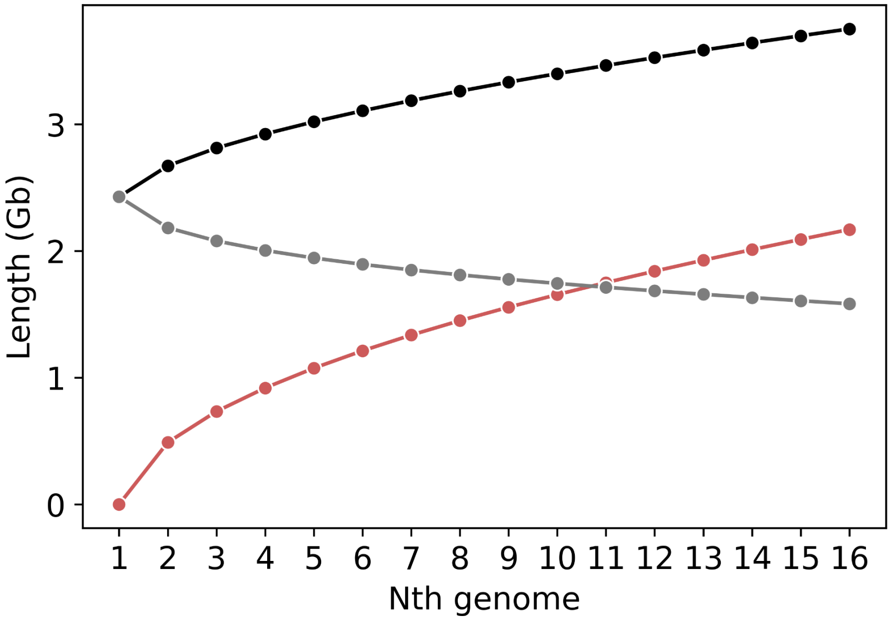


**Extended Data Figure 4:** Growth as a function of increased sampling inferred using panacus for the MC core genome (gray), accessory genome (red), and total pangenome (black).

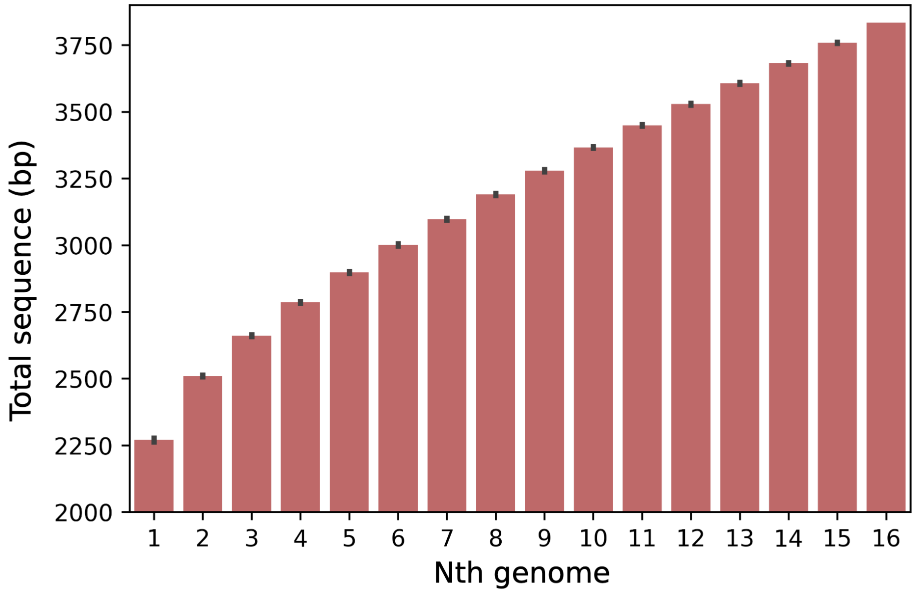


**Extended Data Figure 5:** Bar plot showing the average sequence length of the PGGB pangenome for all permutations of included deer mouse haplotypes. Black lines denote the range of true values across all permutations.

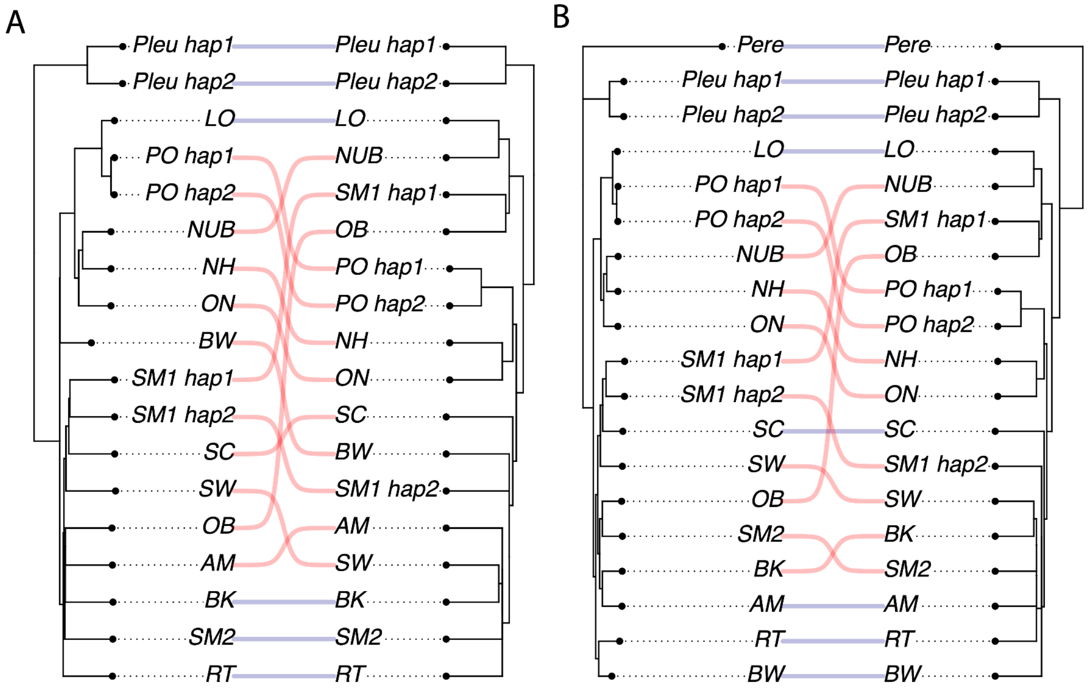


**Extended Data Figure 6:** Topology comparison between (**A**) a bio neighbor-joining (NJ) tree and (**B**) a maximum likelihood (ML) tree constructed from SNPs and a tree constructed from jaccard distances between paths in the PGGB graph (normalized Robinson-Foulds distance = 0.87 and normalized Robinson-Foulds distance = 0.81 for NJ and ML respectively).

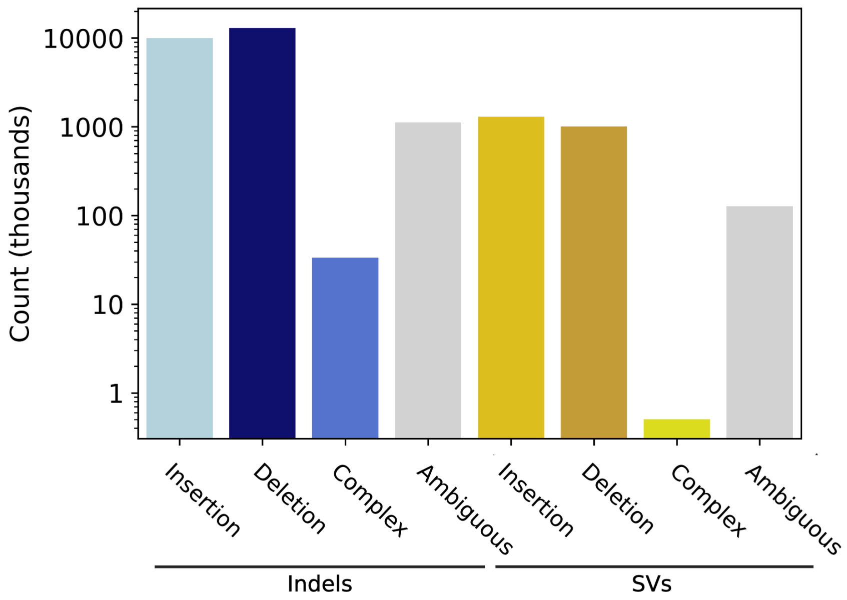


**Extended Data Figure 7:** Variant counts for indel insertions, indel deletions, SV insertions, SV deletions, complex indels and SVs, and ambiguous variants that were multiallelic or could not be confidently polarized.

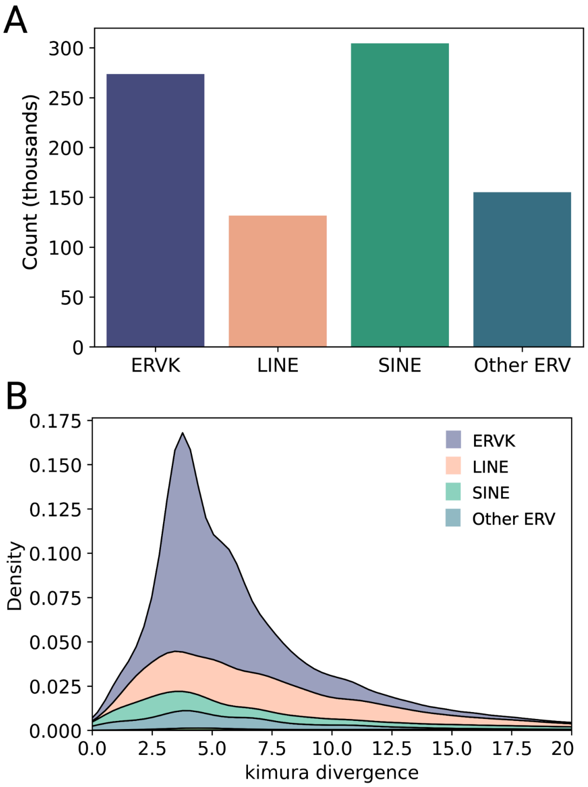


**Extended Data Figure 8:** (**A**) Counts for candidate polymorphic TE insertions across deer mouse haplotypes. (**B**) Distribution of kimura divergence from the consensus values for candidate polymorphic TE copies.

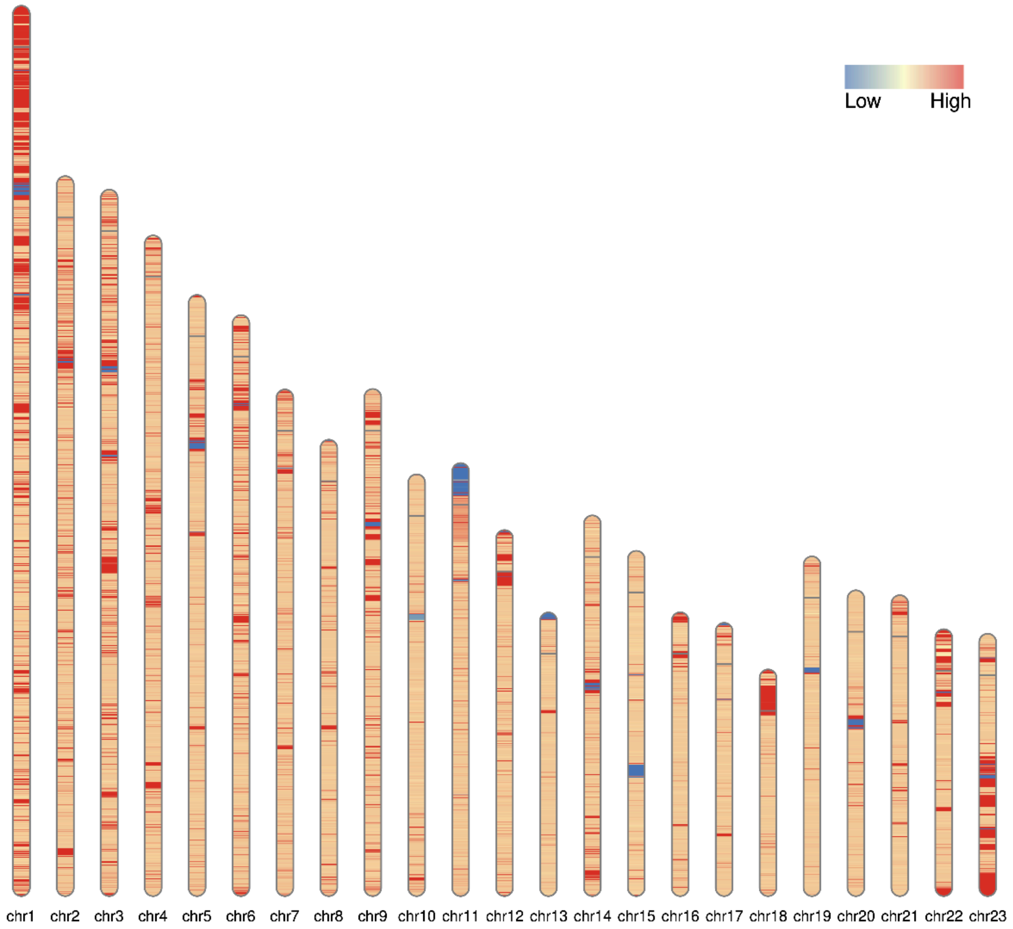


**Extended Data Figure 9:** Ideogram showing node depth distributions across the pangenome projected onto the BW reference. Red regions show high depth and blue regions show low depth.

### Supplemental Table Captions

**Supplementary Table 1. Sample information.** Listed are sample ID, (sub)species, location and approximate GPS coordinates of sampling locations, origin, and source.

**Supplementary Table 2: Assembly statistics.** For each sample, sampleID, subspecies or F1 cross, origin (Lab or wild-caught), sequencing technology, sample barcode, sequencing coverage, assembly software, and assembly_type (primary or haplotype resolved) is listed. For each subsequent assembly we provide the corresponding of number contigs, assembly size (bp), average contig length, scaffold N50, contig N50, contig auN, contig L50, largest contig size, smallest contig size, and BUSCO_S (single), BUSCO_D (duplicate), BUSCO_F (fragmented) BUSCO_I (incomplete), and BUSCO_M (missing) BUSCO scores. We also include QV Scores and depth across scaffolds inferred using Inspector.

**Supplementary Table 3: Repeat content across assemblies.** Percent of the genome occupied by SINEs, LINEs, LTR retrotransposons, DNA transposons and other repeats for each considered assembly.

**Supplementary Table 4: Pangenome graph summary statistics.** Length, node count, edge count, path count, and step count is provided for both PGGB and MC pangenome graphs.

**Supplementary Table 5: Repeat content for accessory and core genomes.** Percent of the genome occupied by SINEs, LINEs, LTR retrotransposons, DNA transposons and other repeats for the core and accessory genomes inferred from the PGGB pangenome graph.

**Supplementary Table 6: Gene copy number variation**. Maximum change in copy number across haplotypes (Max(ΔCN)) for each gene as well as copy respective copy number in each haplotype.

**Supplementary Table 7: CNV gene ontology enrichment results**. For each gene ontology term, we provide the term, core category, term definition, proportion of genes in the study (CNV genes) associated with the term, proportion of genes in the genome associated with the term, p-value, q-value (FDR corrected p-value), and a comma-separated list of CNV genes associated with the term.

**Supplementary Table 8: Allele diversity across loci**. Allele diversity (the number of alleles normalized by the number of haplotypes with calls) for each locus with at least one bubble in the pangene graph. For each locus we provide the genes present and the allele diversity value. Allele diversity could not be calculated for loci with multiple nested bubbles. These loci have NA in the allele diversity column.

**Supplementary Table 9: Key for gene names in Figure 5G,H.** This table lists NCBI gene names and corresponding gene names used in Figure 5 for all genes in Figure 5G and Figure 5H.
